# Identifying cancer cell-state transitions from multimodal single-cell data

**DOI:** 10.64898/2026.03.02.708945

**Authors:** Guido Alessandro Baselli, Alisa Alekseenko, Judit Liaño-Pons, Lefteris Sinanis, Eltjona Rrapaj, Marie Arsenian-Henriksson, Vicent Pelechano

## Abstract

Phenotypic plasticity allows cancer cells to evade therapy, yet the transient nature of state transitions has made their molecular drivers difficult to define. Here, we present a single-cell framework that leverages the temporal delays between mRNA and protein accumulation to directly capture cells undergoing phenotypic switching. We show that differentiation-associated delays between transcript and protein accumulation are detectable in multimodal single-cell data as discordant mRNA and surface-protein levels. Applying this strategy to the K562 leukemia model, which alternates between differentiated CD24- and progenitor-like CD24+ states, we identify transitioning cells and derive a transcriptional signature linking cell-cycle progression and mitochondrial remodeling to plasticity. Genome-wide CRISPR screening confirms key regulators of plasticity, including BCR-ABL1 and mitochondrial homeostasis genes. We summarize the transition-associated program into a score that predicts imatinib response in chronic myeloid leukemia, stratifies survival in acute myeloid leukemia, and retains prognostic value across 31 TCGA tumor types. Spatial transcriptomics reveals localized plasticity hotspots in solid tumors. Together, this framework exposes the molecular basis of cancer plasticity and enables its quantification across tumors.

## Introduction

Dynamic cell-state transitions are increasingly recognized as a fundamental barrier to precision oncology, driving therapeutic resistance and disease progression. While genetic mutations have long been considered the primary source of tumor heterogeneity, non-genetic mechanisms significantly contribute to phenotypic variability, enabling subpopulations of cancer cells to adopt progenitor-like states linked to drug resistance and adverse outcomes^1^. Accordingly, phenotypic plasticity driven by transient epigenetic reprogramming is now considered a hallmark of cancer and is observed across diverse tumor types^2^. Despite its clinical relevance, the molecular programs governing these transitions remain poorly characterized because cells in active transition are transient and difficult to capture. Addressing this gap is essential to understand how oncogenic signaling and gene expression remodeling cooperate to generate therapy-resistant states and to develop strategies to target them.

Most single-cell methods provide a static snapshot of a cell population. State transitions can be inferred from single-modality data such as single-cell transcriptomics (scRNA-seq) and chromatin accessibility profiling (scATAC-seq)^3, 4^, but such inference can be confounded by multiple sources of variation (e.g., cell cycle) and transition directionality is often ambiguous^5^. Dynamic approaches such as RNA velocity can provide more direct estimates of cell trajectories^6^. However, RNA velocity relies on sparse unspliced (intronic) read counts and on kinetic assumptions that may be violated in specific biological contexts^7^. Although recent velocity algorithms relax some of these assumptions^8^, integrating orthogonal data layers offers a complementary strategy to directly identify cells undergoing state transitions.

Here, we present a single-cell framework that leverages the temporal delay between mRNA and protein expression dynamics to directly identify cells undergoing state transitions and define their transcriptional signature. We apply this strategy to the chronic myeloid leukemia (CML) cell line K562^9, 10^, which differentiates along the erythroid lineage and reversibly switches between a differentiated CD24-phenotype and a progenitor-like, drug-resistant CD24+ state^11^. By integrating surface-protein quantification with single-cell transcriptomics, we capture cells transitioning between these states and define a transcriptional signature of cell plasticity. Furthermore, we show that changes in canonical markers mRNA levels precede corresponding protein-level changes during monocyte differentiation, supporting the generalizability of this framework.

In the K562 cells, this framework uncovers a mechanistic link between cell-cycle progression, mitochondrial remodeling, and phenotypic switching, which we validate through a genome-wide CRISPR knockout screen. To evaluate the generalizability of these findings beyond the K562 model, we assessed the association between the plasticity-associated signature and clinical outcomes across cancer cohorts. Summarized as a plasticity score, the signature predicts drug response in CML and overall survival across three independent acute myeloid leukemia (AML) cohorts. Extending these associations beyond hematologic malignancies, the plasticity score also predicts outcome across 31 non-hematological tumor types in TCGA and identifies intra-tumoral high-plasticity hotspots in spatial transcriptomics data from hepatocellular carcinoma and clear-cell renal cell carcinoma.

Overall, our results underscore the utility of protein-mRNA dynamics for identifying and studying phenotypic state transitions and establish a framework that is broadly applicable to multimodal single-cell datasets. We highlight plasticity-associated transcriptional programs that are shared across tumors and may be leveraged as biomarkers and therapeutic targets in oncology.

## Results

### K562 cells exhibit reversible phenotypic states

K562 cells exist in a dynamic equilibrium between differentiated (CD24-) and progenitor-like (CD24+) states^11^. To confirm this reversibility, we isolated CD24+ and CD24-cells by immunomagnetic sorting and monitored CD24 surface expression over time. Within six days, both populations converged to overlapping immunophenotypes (Figure 1A), demonstrating rapid and reversible switching. We investigated the transcriptional differences using single-cell RNA sequencing combined with surface antigen barcoding to quantify protein levels (Figure 1B, Figure S1 and Supplementary Materials and Methods).

**Figure 1.**
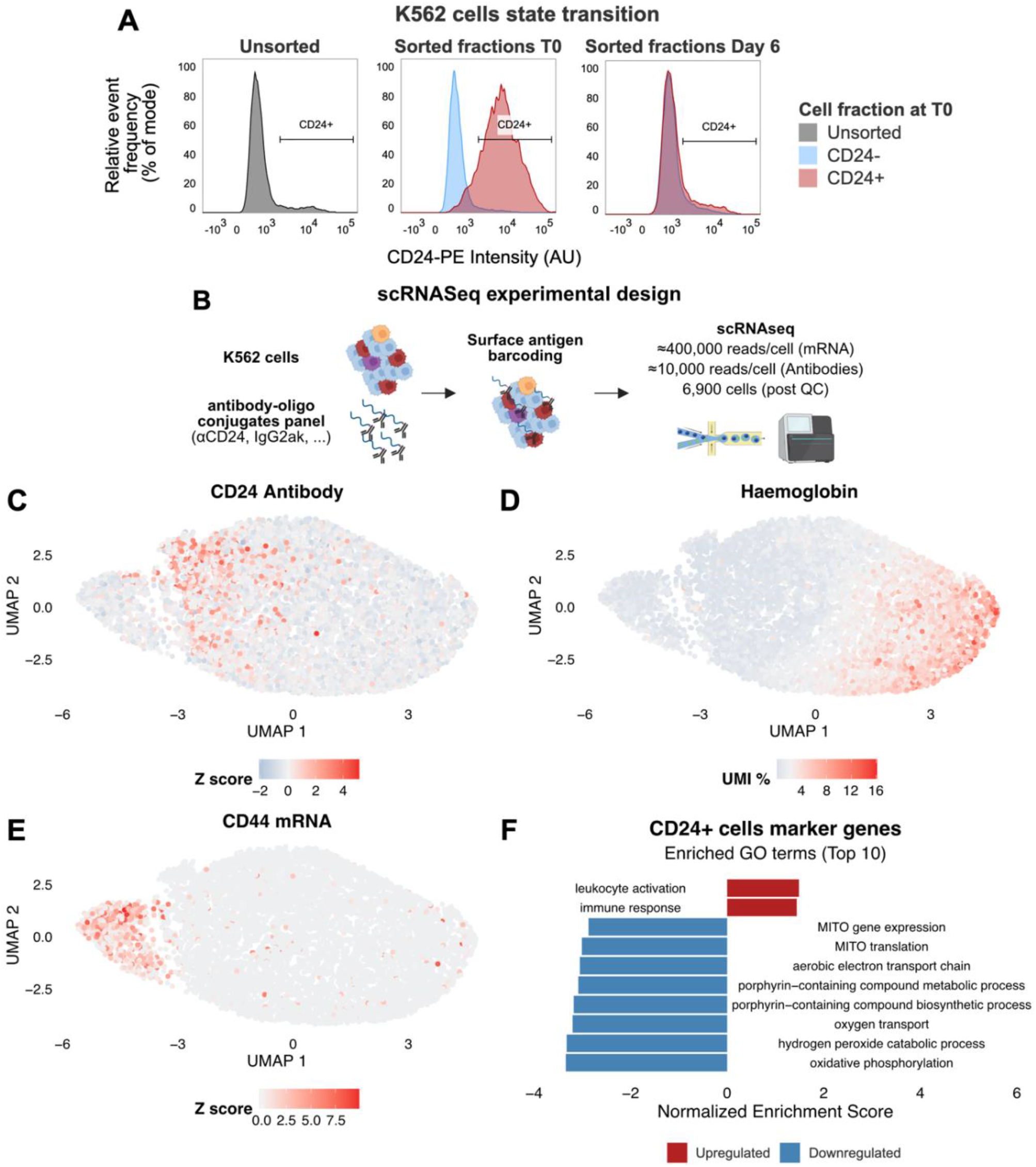
Composition of the K562 cell line and characterization of the CD24+ phenotype. **A)** Frequency histograms of CD24+ and CD24-K562 cells isolated by immunomagnetic sorting (MACS) and analyzed by flow cytometry at T0 and six days post sorting. **B)** scRNA-seq experimental workflow. K562 cells were labeled with surface antigen antibodies and analyzed by scRNA-seq to simultaneously quantify surface protein expression and transcriptomic profiles within the same cells. **C-E)** Uniform Manifold Approximation and Projection (UMAP) of scRNA-seq data from K562 cells, colored by CD24 protein abundance (C), hemoglobin gene expression measured as the per-cell proportion of hemoglobin UMIs relative to the total UMI count (D), or CD44 expression (E). **F)** Top ten Gene Ontology (GO) terms enriched in genes upregulated or downregulated in CD24+ versus CD24-cells as assessed by Gene Set Enrichment Analysis (GSEA).

Uniform Manifold Approximation and Projection (UMAP) revealed a heterogeneous main cluster including cells with high CD24 protein levels and cells with high hemoglobin mRNA expression (Figures 1C-D), consistent with a less differentiated phenotype in CD24+ cells. Bone marrow reference mapping confirmed the erythroblast-like identity of K562 cells (Figure S2A) and showed that CD24+ cells are enriched in earlier differentiation stages (Figure S2B). We also identified a smaller cluster of CD44+ cells, possibly representing an alternative differentiation trajectory (Figure 1E).

Differential gene expression analysis revealed that CD24+ cells downregulate genes involved in erythroid differentiation and mitochondrial metabolism, while upregulating genes associated with leukocyte proliferation and activation (Figure 1F and Tables S1-2).

We validated these findings using an independent K562 scRNA-seq dataset^12^, where we found an inverse correlation between CD24 mRNA levels and expression of mitochondrial metabolism-related genes (Figure S2C). Furthermore, sorted CD24+ K562 cells exhibited lower baseline oxygen consumption rate (OCR) as assessed by extracellular flux analysis (Figure S2D-E). Conversely, CD44+ cells overexpressed plasmablast-, megakaryocyte-, and stemness-associated markers (Tables S3-4) consistent with a diverging differentiation branch. Together, these findings reveal that K562 cells exist in a dynamic equilibrium between a metabolically active erythroid-like state and a progenitor-like, metabolically quiescent CD24+ state, with CD44+ cells representing a diverging trajectory.

### Protein-mRNA discordance identifies transitioning cells in multimodal single-cell data

To define the molecular features of cells actively switching states, we leveraged the temporal delay between mRNA and protein expression as a proxy for transition. Because transcription precedes protein accumulation and mRNA generally decays faster than its corresponding protein^13^, cells undergoing transition are expected to display discordant CD24 transcript and protein levels (Figure 2A).

**Figure 2.**
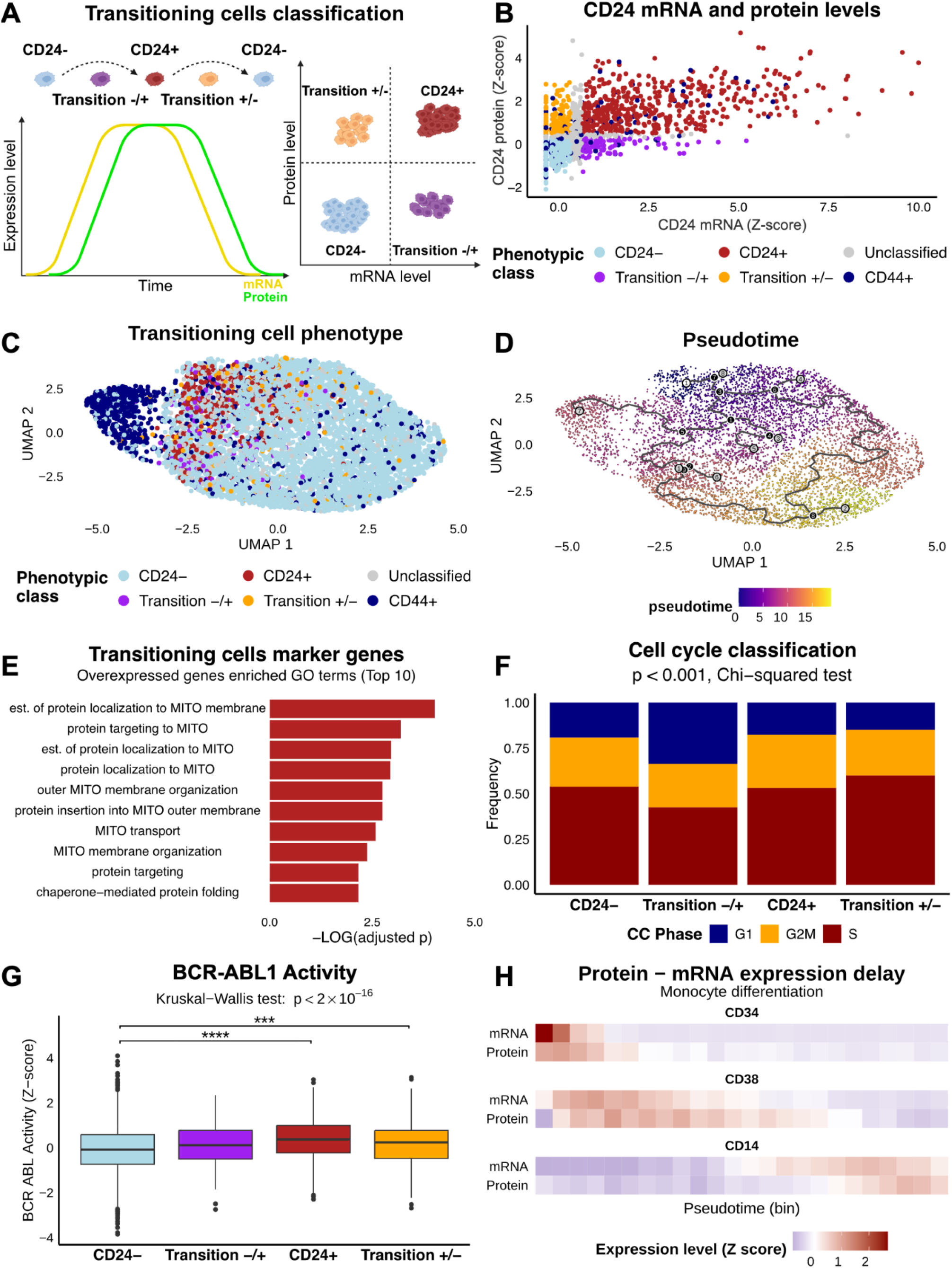
Protein-mRNA discordance identifies transitioning cells in CITE-Seq data. **A)** Rationale for identifying transitioning K562 cells based on CD24 mRNA and protein levels. In CD24-cells beginning to express CD24, mRNA accumulates first, followed by protein accumulation. Conversely, in CD24+ cells downregulating CD24, mRNA levels decrease before protein levels are lost. Therefore, cells undergoing active transitions are expected to display discordant mRNA and protein levels. As shown in the right panel, cells in the -/+ transition are expected to show high mRNA levels with low protein expression, whereas cells in the +/-transition are expected to exhibit low mRNA levels while retaining high protein levels. **B)** CD24 protein and mRNA levels in K562 cells colored by phenotypic class: CD24+, CD24-, Transition -/+, Transition +/-, and CD44+. **C)** Uniform Manifold Approximation and Projection (UMAP) of scRNA-seq data from K562 cells colored by phenotypic class. **D)** Pseudotime trajectory analysis in K562 cells. Black lines represent inferred cell-state trajectories, with numbered circles indicating the root point (white), branch points (black), and terminal states (grey). **E)** Top ten Gene Ontology (GO) terms enriched in genes overexpressed in transitioning cells, assessed by overrepresentation analysis. **F)** Cell-cycle phase distribution across CD24+, CD24-, and transitioning cells. A chi-squared test was performed to assess variation in cell proportions across phenotypic states. **G)** BCR-ABL1 signaling activity scores by phenotypic class. Statistical analysis was performed using the Kruskal-Wallis test followed by Dunn’s post-hoc test with Holm correction for multiple comparisons. **H)** RNA and protein expression levels of canonical differentiation markers CD34, CD38, and CD14 along pseudotime in differentiating monocytes from public bone marrow CITE-seq data.

To capture this discordance, we modeled the relationship between CD24 mRNA and protein abundance (Figure S3A), predicted expected protein levels from transcript counts, and assigned cells to predicted CD24+ or CD24-states using the same thresholds as for the observed protein. Cells with discordant predicted and observed CD24 states were classified as transitioning (Figure 2B). To avoid confounding from diverging lineages, CD44+ cells were excluded from the analysis.

Cells in -/+ and +/-transition were rare, representing 1.6% and 3.4% of cells, respectively (Figure S3B), and showed preferential localization in UMAP space (Figure 2C). Consistently, each transitioning population differed from the other phenotypic classes along at least one UMAP component (Figure S3C-D), suggesting distinct transcriptional states.

Trajectory analysis resolved two branches connecting CD24+ and CD24-cells and further supported CD44+ cells as a diverging branch (Figure 2D). Pseudotime ordering was consistent with the inferred differentiation status of K562 cells (Figure S3E). As expected, both transitioning populations displayed pseudotime values higher than CD24+ cells but lower than CD24-cells (Figure S3F), supporting their transitional nature.

To identify pathways driving state transitions, we performed pseudobulk differential expression analysis (Table S5). Transitioning cells (-/+ and +/-) overexpressed genes involved in mitochondrial homeostasis (Figure 2E and Table S6), suggesting that mitochondrial remodeling facilitates shifts between the metabolically distinct phenotypes.

We next examined whether state transitions correlate with cell-cycle dynamics driven by the BCR-ABL1 fusion oncogene. Cell-cycle analysis showed a significant association between transitions and cel-cycle stage (Figure 2F). Cells transitioning to CD24+ were enriched in G1 phase and less frequently in S phase, whereas those reverting to CD24-were more frequent in S phase. Consistently, BCR-ABL1 signaling activity varied across states, peaking in CD24+ cells and reaching a minimum in CD24-cells (Figure 2G).

To test whether this approach generalizes beyond K562 cells, we reanalyzed public bone marrow CITE-seq data^14^ and focused on the differentiation trajectory from CD34+CD38-hematopoietic stem cells toward CD14+ classical monocytes, using pseudotime to model progression through differentiation stages (Figure S4A-C). Along pseudotime, canonical differentiation markers showed delayed protein-level changes relative to mRNA-level changes (Figure 2H).

Collectively, these observations suggest that protein-mRNA discordance in differentiation markers can identify transitional cell states. In K562 cells, this framework highlights a link between BCR-ABL1-driven cell-cycle progression, metabolic rewiring, and phenotypic switching between the CD24+ and CD24-states.

### Mitochondria homeostasis and oncogenic signaling govern plasticity

To identify genetic determinants of phenotypic transition, we performed a genome-wide CRISPR knockout screen using a two-step sorting strategy to enrich for transitioning cells (Figure 3A). To validate the screening platform, we first compared CD24+ and CD24-cells after the first sorting step to identify sgRNAs associated with the CD24+ phenotype. As detailed in the Supplementary Results, sgRNAs targeting genes required for CD24 biosynthesis were depleted from CD24+ cells (Figure S5, Tables S7-8).

**Figure 3.**
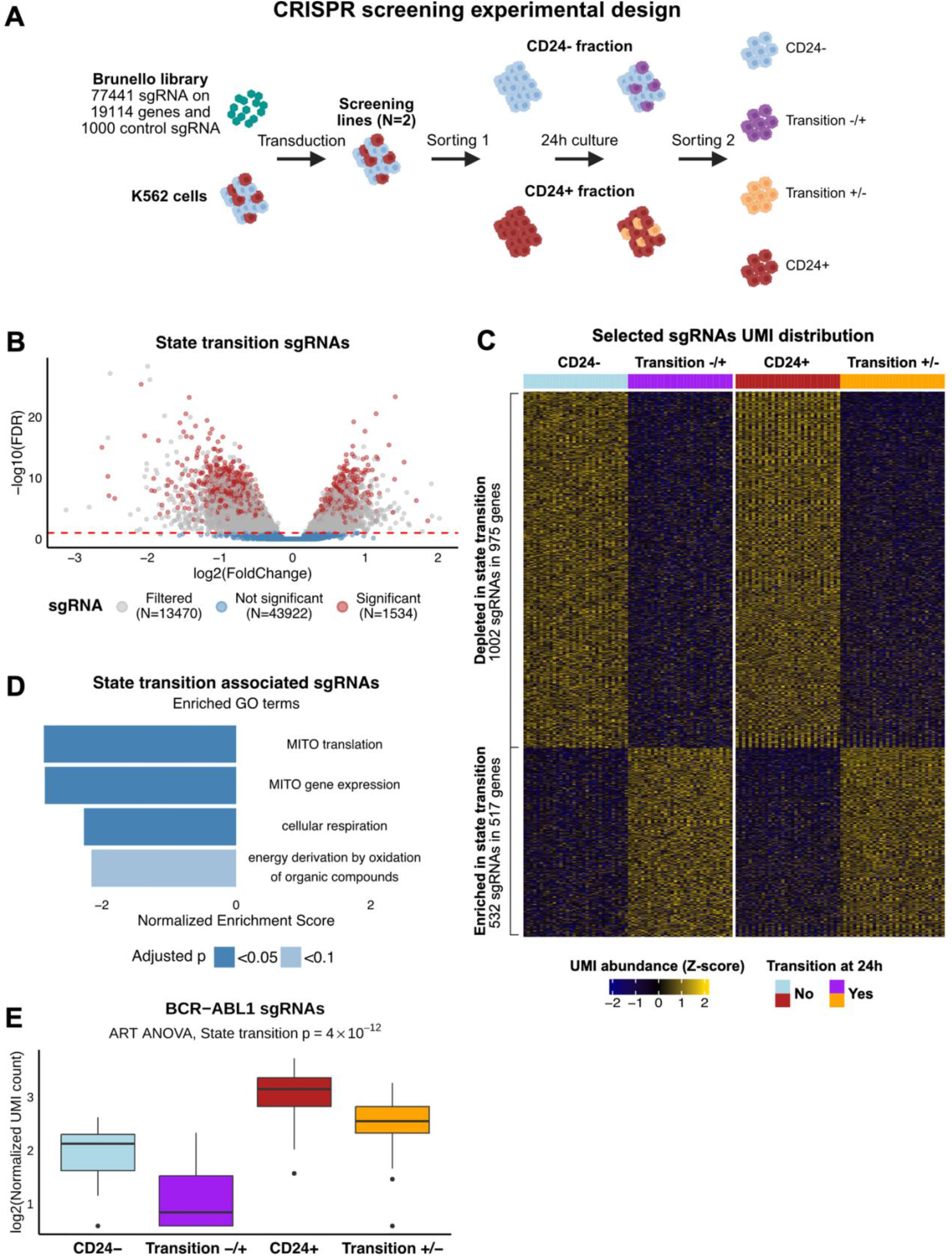
Genetic drivers of state transition in K562 cells. **A)** CRISPR screening experimental design. CD24+ and CD24-cells were isolated by immunomagnetic sorting (MACS) and cultured for 24 h. Cells were then sorted again to isolate stable CD24+ and CD24-cells, as well as cells that underwent a -/+ or +/-state transition. **B)** Volcano plot showing differential abundance of the sgRNAs in transitioning versus stable cells. Statistically significant sgRNAs selected for downstream analysis, as described in the Methods section, are marked in red. **C)** Heatmap displaying the distribution of abundance values for the selected single guide RNA (sgRNAs) in the two transitioning populations compared with their stable counterparts. **D)** Gene Ontology (GO) terms enriched in genes targeted by sgRNAs depleted in transitioning cells, as assessed by Gene Set Enrichment Analysis (GSEA). **E)** Abundance distributions of sgRNAs (N=3) targeting the BCR-ABL1 fusion gene across the cell states. Statistical analysis was performed by logistic regression.

We then considered sgRNAs associated with state transition independently of transition direction (Figures 3B-C, Table S9). Among these, sgRNAs targeting mitochondrial gene expression and oxidative phosphorylation pathways were depleted in transitioning populations (Figure 3D, Table S10), supporting mitochondrial homeostasis as a key regulator of plasticity. Among plasticity-associated sgRNAs (Table S9), we identified guides targeting ABL1 and BCR-ABL1 effectors, including STAT3 and HRAS. Comprehensive analysis of all sgRNAs targeting BCR-ABL1 revealed significant depletion of BCR-ABL1 knockout events in both transitioning populations compared with their stable counterparts (Figure 3E).

Together, these findings indicate that BCR-ABL1 signaling, and mitochondrial remodeling contribute to phenotypic switching, reinforcing the mechanistic link observed in our transcriptomic analyses.

### Cell plasticity signature predicts patient outcome across tumor types

Having functionally validated state-transition regulators in the K562 model, we next assessed the clinical relevance of our findings in patient data. CD24+ cells have previously been linked to drug resistance in CML models^11^. Building on this, we hypothesized that the transcriptional plasticity signature defined in our single-cell analyses captures clinically relevant features of therapy resistance. To test this, we assessed the association between the plasticity signature and therapeutic response in CML. We reanalyzed gene-expression profiles collected at diagnosis from 96 CML patients initiating imatinib treatment^15^ and summarized the expression of transition-associated genes into a single plasticity score per patient (see Supplementary Results and Figure S6 for score definition and validation). Patients with high plasticity scores showed significantly reduced early molecular response at three months (EMR; Figure 4A), consistent with reduced imatinib sensitivity.

**Figure 4.**
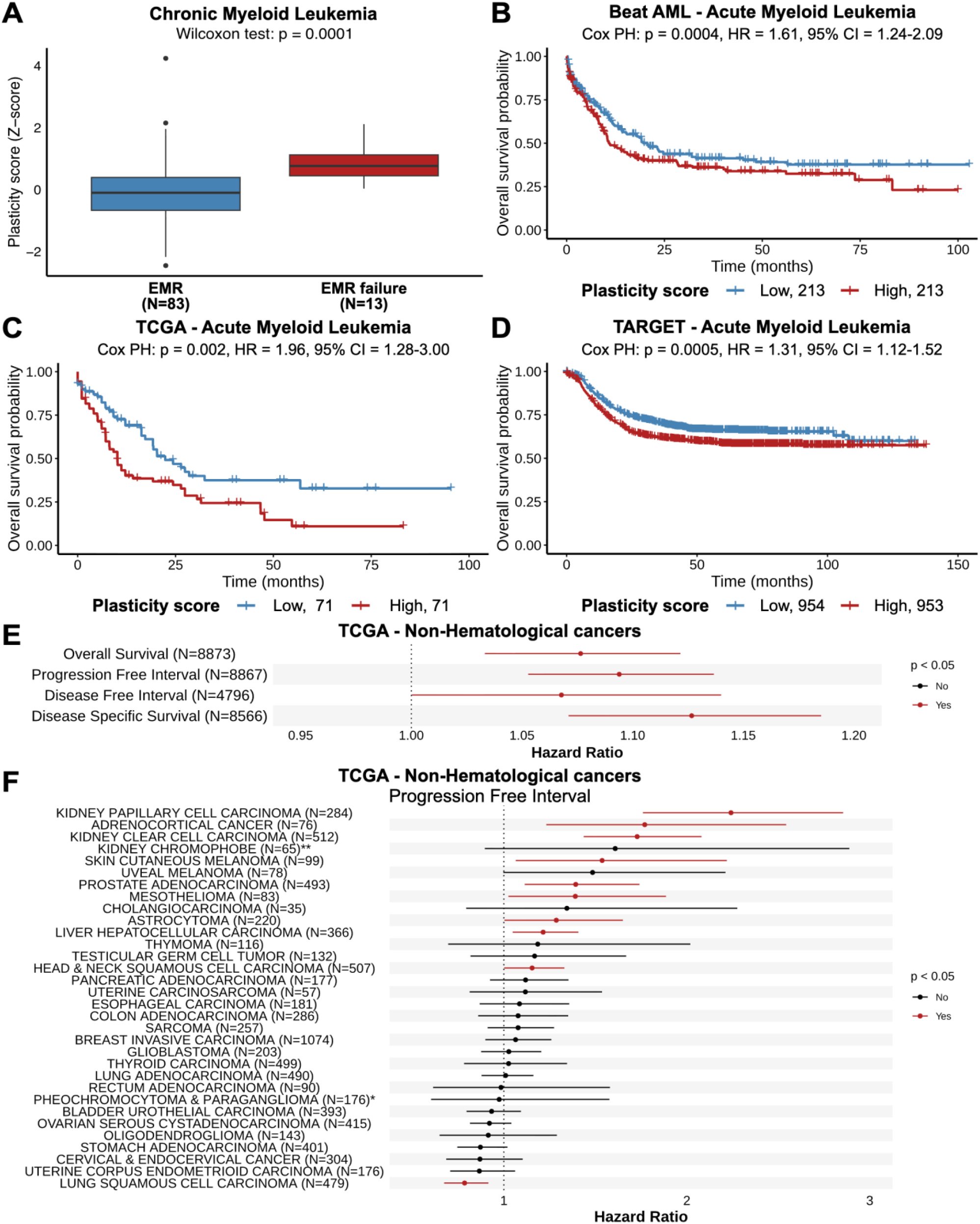
Plasticity score clinical associations. **A)** Plasticity score distributions in a cohort of 96 CML patients stratified by early molecular response (EMR) failure. **B-D)** Kaplan-Meier survival curves in the Beat AML (B, N = 426), TCGA-LAML (C, N = 142), and TARGET-AML (D, N = 1,907) cohorts stratified by plasticity score group. **E-F)** Forest plots showing associations between plasticity score and clinical outcomes across TCGA non-hematological cancers: pan-non-hematological-tumors analysis (E) and progression-free-interval analyses in individual tumor types (F). Differential analysis in A was performed using the Wilcoxon test. Associations with outcomes (B-F) were assessed using age-adjusted Cox proportional hazards regression. HR, hazard ratio; CI, confidence interval. In F, * and ** denote cohorts not recommended or recommended with caution for this outcome, respectively, according to TCGA guidelines.

Although our mechanistic data indicate that BCR-ABL1 signaling drives plasticity in K562 cells, we reasoned that the underlying principle (cell-cycle-linked metabolic rewiring) could be conserved across malignancies. We therefore assessed whether the plasticity score predicted overall survival in three independent AML cohorts: Beat AML^16^, TCGA-LAML^17^, and the pediatric TARGET-AML group^18^ (Table S11). Higher plasticity scores were associated with worse overall survival after adjustment for age in each cohort and in the pooled analysis across all AML patients (Figure 4B-D; Table S12).

To explore relevance in solid tumors, we extended the scoring approach to 31 non-hematological cancer types from TCGA^19^ (Table S13). The plasticity score was associated with worse overall survival, progression-free interval, disease-free interval, and disease-specific survival after adjusting for age and stratifying by tumor type (Figure 4E; Table S14). These associations remained robust in sensitivity analyses restricted to cohorts with adequate event counts, except for disease-free interval (Figure S7A; Table S15). Analyses within individual cancer types confirmed higher plasticity score associations with shorter progression-free interval (Figure 4F) and overall survival (Figure S7B; Table S16) in clinically relevant solid tumors, including kidney clear cell (ccRCC) and hepatocellular carcinoma (HCC).

Taken together, these results show that the transcriptional program underlying cell-state plasticity has broad prognostic significance across myeloid leukemias and solid tumors. The ability of the plasticity score to stratify outcomes across cancer lineages suggests a conserved biological basis and highlights phenotypic plasticity as a clinically relevant dimension of tumor behavior.

### Plasticity signature identifies proliferative hotspots in solid tumors

Because solid tumors display marked spatial metabolic heterogeneity, we asked whether plasticity localizes to specific intratumoral niches. We re-analyzed public spatial transcriptomic profiles from two HCC sections from two different patients with distinct histopathological features^20^ (Figure 5A, Figure S8A-B, and Supplementary Results). In these data, per-spot plasticity scores correlated positively with signature scores related to proliferation, organelle division, and oxidative metabolism, and negatively with fatty acid metabolism programs (Figure 5B, Table S17, and Supplementary Results). Spatial mapping revealed high-plasticity spots in peri-fibrotic or highly proliferative areas, which also exhibited elevated proliferation, oxidative phosphorylation, and organelle fission scores (Figures 5C-F and S8C-F).

**Figure 5.**
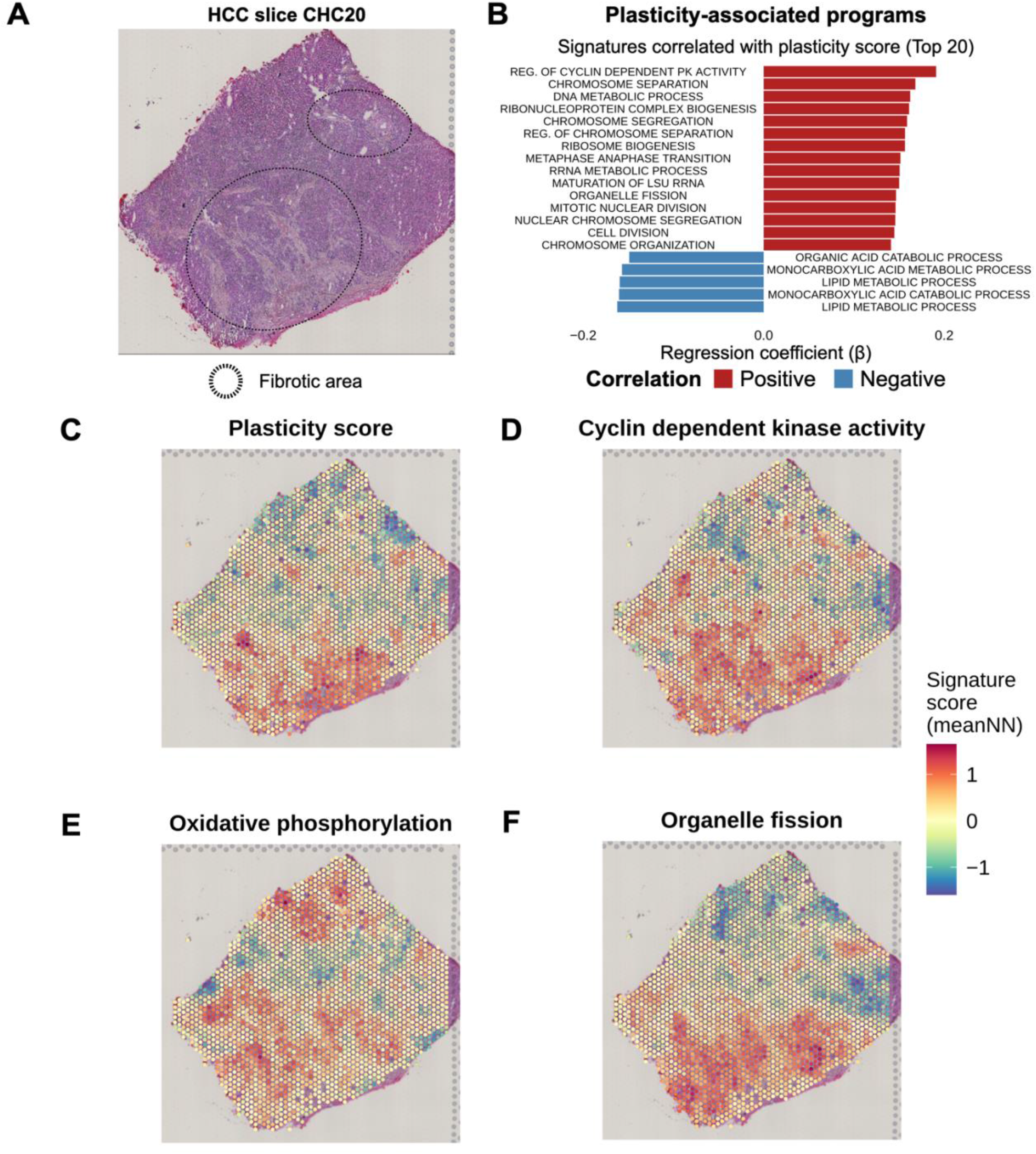
Plasticity score in HCC spatial transcriptomics. **A)** H&E staining showing the morphology of the CHC20 slice. Fibrotic regions are highlighted; no clear non-tumoral areas were detected in this section. **B)** Top 20 GO Biological Process signatures correlated with spot-level plasticity scores. **C-F)** Spatial distribution of the Plasticity score in CHC20 HCC slice (C), Cyclin-dependent protein kinase activity (D), Oxidative phosphorylation (E), and Organelle fission (F). For visualization, signature scores were smoothed using a mean nearest-neighbor approach (k = 6).

Analogous results were observed in a ccRCC dataset comprising 12 sections from 12 patients^21^, as described in the Supplementary Results (Figure S9 and Table S18).

Together, these analyses show that the plasticity signature resolves localized state-transition hotspots within solid tumors and highlights the conservation of this transcriptional program across HCC and ccRCC.

## Discussion

The ability of cancer cells to reversibly transition between phenotypic states is increasingly recognized as a major contributor to therapeutic resistance and disease progression^1^. In the K562 model of chronic myeloid leukemia, earlier work established a dynamic equilibrium between a drug-sensitive CD24-state and a progenitor-like CD24+ state^11^. Here, we used this system to investigate the molecular features of cells undergoing phenotypic transition and to dissect the mechanisms that govern this process.

We first confirmed the reversible nature of the progenitor-like CD24+ state^11^ and showed that CD24+ cells rely less on oxidative phosphorylation than their CD24-counterparts.

We then leveraged the temporal delay between mRNA and protein accumulation to identify cells in active transition using single-cell multimodal data. This strategy is based on the observation that protein turnover generally occurs on a longer timescale than mRNA turnover^13^. We showed that protein-level changes lag mRNA-level changes for canonical markers along monocyte differentiation. Thus, this approach can be readily applied to multimodal single-cell datasets.

In K562 cells, this framework allowed us to define a transcriptional signature characterizing state transition, irrespective of transition direction. We found that transitioning cells overexpress genes involved in mitochondrial homeostasis, suggesting a link between state transitions and metabolic reprogramming, consistent with emerging crosstalk between metabolism and epigenetic regulation^22^. Further analyses revealed altered cell-cycle phase distributions in transitioning populations and substantial variation in BCR-ABL1 signaling activity across phenotypic classes. Together, these observations support a model in which aberrant BCR-ABL1-driven proliferation is coupled to metabolic rewiring that facilitates phenotypic switching, consistent with evidence linking oncogene-driven growth to metabolic adaptation^23, 24^ and with reports of metabolic plasticity in myeloid malignancies^25-27^.

To functionally validate these mechanisms, we performed a genome-wide CRISPR knockout screen within a sequential sorting framework designed to enrich for lineages with altered state-transition kinetics. This analysis confirmed that mitochondrial homeostasis and oxidative metabolism are key regulators of plasticity and highlighted a role for BCR-ABL1 signaling in promoting state transitions. Collectively, the CRISPR results aligned with our scRNA-seq observations, providing orthogonal validation of our single-cell-based analytical framework and supporting the link among cell-cycle, metabolic reprogramming, and state transition.

Given the established link between CD24+ cells and drug resistance^11^, we hypothesized that plasticity influences therapeutic response. Consistent with this, the plasticity score derived from our single-cell analyses predicted early imatinib response in CML patients.

Although BCR-ABL1 drives plasticity in our CML model, the broader principle that oncogene-driven proliferative signaling is coupled to metabolic reprogramming and phenotypic plasticity is likely more general^23, 24^. In line with this, the plasticity score predicted survival across three independent AML cohorts^16, 18, 28^, consistent with the established importance of metabolic programs in AML progression and resistance^25, 26^.

To assess generalizability beyond hematological cancers, we evaluated the plasticity score across 31 solid tumor types in TCGA^19^. The score was consistently associated with poorer outcomes across tumor types, including overall survival and progression-free interval. Notably, analyses within individual tumor types revealed strong associations in clinically relevant cancers such as ccRCC and HCC.

Moreover, spatial transcriptomic analyses of these tumors showed that high plasticity regions localized to discrete intratumoral domains enriched for proliferation, mitochondrial activity, and organelle fission. These spatial patterns mirror the mechanistic features observed in K562 cells and raise the possibility that mitochondrial division contributes to proliferation-linked metabolic remodeling during state transitions.

Although the plasticity score was broadly associated with poorer outcomes across cancer types, we also detected sporadic opposite associations in individual tumors as lung squamous cell carcinoma. This suggests that the biological significance of the transcriptional program identified here may differ across tumor lineages, and further studies will be required to investigate plasticity across heterogeneous contexts.

In summary, joint single-cell measurement of mRNA and protein enabled us to identify cells undergoing state transitions and to characterize their transcriptional programs. This approach defined a transition-associated transcriptional signature and linked plasticity to cell-cycle progression and metabolic reprogramming. The resulting plasticity score predicts therapeutic response in CML and prognosis in AML and diverse solid tumors and reveals discrete plasticity hotspots in spatial transcriptomic data from HCC and ccRCC. Together, these findings suggest shared mechanisms linking proliferation, metabolism, and cell-state transitions across cancer types, and provide a broadly applicable framework for quantifying plasticity from single-cell multimodal data. Future studies will be needed to determine whether targeting metabolic rewiring or state-transition programs can improve patient outcomes.

## Materials and Methods

Procedures used for flow cytometry analysis, reanalysis of the validation scRNASeq dataset (GSE855348) ^12^, extracellular flux analysis, reanalysis of public bone-marrow CITE-seq data^14^ (GSE245108) and spatial transcriptomics data^20, 21^ (GSE245908 and GSE175540) are described in the Supplementary Materials and Methods.

### Cell culture

K562 cells (ATCC, CCL-243) were cultured at 37°C in a humidified incubator with 5% CO_2_ in RPMI-1640 medium with L-glutamine and sodium bicarbonate (Sigma-Aldrich, R8758) supplemented with 10% fetal bovine serum (Sigma-Aldrich, F7524) and 100 U/ml penicillin-streptomycin (Sigma-Aldrich, P4333). Cells were passaged every three days by seeding 10^5^ cells/ml in fresh medium.

### 10x scRNA-seq and feature barcoding

Detailed library-preparation protocols and analysis procedures are provided in the Supplementary Materials and Methods.

Briefly, three independent K562 samples were stained with a panel of oligo-conjugated antibodies (BioLegend, TotalSeq B), including anti-CD24, anti-CD34, anti-CD38, two isotype controls (IgG2aκ and IgG1κ), and a unique cell hashing antibody (HTO) for each technical replicate. Libraries were prepared using the Single Cell 3^′^ Reagent Kits v3.1 with Feature Barcoding technology (10x Genomics) and sequenced on NovaSeq 6000 and NovaSeq X platforms to a depth of ≈4*10^5^ reads per cell for the mRNA library and 10^4^ reads per cell for the antibody library. Read alignment and UMI counting were performed using CellRanger v7.1.0 (10x Genomics) and the GRCh38-2020-A reference.

Data quality control and analysis were performed in R v4.4.0 using Seurat v5.3.0^29^. CD24 antibody counts were normalized using the matched isotype control (IgG2aκ), and normalized CD24 protein levels were used to classify CD24- and CD24+ cells (Z-score < 0.3 and > 0.5, respectively; Figure S1F). CD44+ classification was based on CD44 mRNA detection, defined as UMI count > 0.

For downstream transition-state analyses, differential expression in transitioning cells was assessed by pseudobulk DESeq2 analysis across phenotypic classes and biological replicates, followed by pathway enrichment using ClusterProfiler. Continuous variables were compared across groups using the Kruskal-Wallis test followed by Dunn’s post-hoc test with Holm correction for multiple comparisons. Associations between categorical variables were assessed using the chi-squared test.

### Genome-wide CRISPR-Cas9 knockout screening

Full line-generation, sorting, library-preparation, and data-analysis protocols are provided in the Supplementary Materials and Methods. Briefly, two independent CRISPR-KO screening lines were generated using the genome-wide Brunello sgRNA library^30^ and a UMI-based strategy to track individual transduction events^31^. To enrich for transitioning cells, we used a sequential sorting procedure. Cells were first sorted into CD24+ and CD24-populations using the CELLection Pan Mouse IgG Kit (Thermo Fisher, 11531D) and an anti-CD24 antibody (BD 555426, clone ML5). Both fractions were cultured for 24 h and sorted again, yielding CD24-, Transition -/+, CD24+, and Transition +/-populations (Figure 3A). Samples for sequencing were collected at each step: unsorted cells, first-sort fractions, and second-sort fractions. Libraries were prepared as previously described^31^ and sequenced on a NextSeq 2000 (Illumina). CRISPR-screening data were analyzed using an internal-replicate framework based on lineage-tracking UMIs^31^. Differential sgRNA abundance was assessed using DESeq2, adjusting for biological replicate and, where appropriate, initial CD24 state. Hits were filtered to retain sgRNAs with concordant effects in -/+ and +/-transitions and to remove sgRNAs targeting genes with discordant knockout effects. BCR-ABL1 was analyzed by considering all sgRNAs targeting the BCR-ABL1 locus using a two-way aligned rank transform (ART) ANOVA framework accounting for state transition and initial cell state.

### Clinical cohort plasticity scoring

Clinical cohort selection, filtering, plasticity score calculation, and analysis procedures are described in detail in the Supplementary Materials and Methods.

Briefly, CML gene-expression data from 96 chronic-phase patients profiled at diagnosis were obtained from GEO (GSE130404)^15^. AML analyses included patients from the Beat AML^16^ (N = 426), TCGA-LAML^17^ (N = 142), and TARGET-AML^18^ (N = 1,907) cohorts. Gene-expression and clinical data for these AML cohorts were obtained via cBioPortal^32^ or the GDC data portal v43 ^33^. For pan-cancer analyses, normalized gene expression data and clinical data from 8,873 patients with primary non-hematological tumors from TCGA^19, 28^ were obtained from UCSC Xena^34^.

Plasticity scores were calculated by rank-deviation. Briefly, genes were ranked by expression within each sample, compared with their average rank across the cohort, signed according to their direction in the transition-associated signature, summed, and Z-score normalized. For cohorts with multiple sampling sites or tumor types, score calculation and normalization were performed within sampling site or tumor type. Associations with binary outcomes were assessed using Wilcoxon rank-sum tests, whereas survival associations were evaluated using age-adjusted Cox proportional hazards models, with tumor type included as a stratification variable where indicated.

## Supporting information

Supplementary results, supplementary methods, Supplementary Figures, Supplementary Figures and Tables legends

## Ethics

Reanalysis of published clinical data was approved by the Swedish ethics review authority (Etikprövningsmyndigheten reg. 2025-02231-01).

## Acknowledgements

We thank Pareja-Sanchez Y and Brinkman EK for their contribution to the early stages of the project. We also thank all Pelechano (Karolinska Institutet, SciLifeLab), Kutter (Karolinska Institutet, SciLifeLab), and Friedlander (Stockholm University, SciLifeLab) labs members for their contribution to the project through scientific discussions. We acknowledge the Biomedicum Flow Cytometry Core Facility (Karolinska Institutet), supported by KI/SLL, for providing cell sorting and analysis services and technical expertise. We thank the CRISPR Functional Genomics Unit (CFG) at Karolinska Institutet funded by SciLifeLab for their contribution to the CRISPR screening experimental design and screening line generation. The authors acknowledge support from the National Genomics Infrastructure in Stockholm, funded by Science for Life Laboratory, the Knut and Alice Wallenberg Foundation, and the Swedish Research Council, as well as SNIC/Uppsala Multidisciplinary Center for Advanced Computational Science, for assistance with scRNA-seq library preparation, massively parallel sequencing, and access to the UPPMAX computational infrastructure. The computations were enabled by resources provided by the National Academic Infrastructure for Supercomputing in Sweden (NAISS), partially funded by the Swedish Research Council through grant agreement no. 2022-06725. Figures 1B, 2A, and 3A were created with BioRender. Microsoft copilot was used to polish the text, authors review those edits and take full responsibility for the manuscript.

## Authors Contribution

G.A.B., V.P. and A.A., conceived the project. G.A.B. led the experimental design and conducted the bulk of wet lab procedures and computational analyses. A.A., J.L.-P., L.S. and E.R. contributed to the experimental procedures and design. M.A.-H. contributed to experimental design and supervision. V.P. curated the study design and supervised the study.

G.A.B. wrote the initial manuscript with the help of V.P. All authors contributed to the manuscript and approved the final version.

## Data Availability

Upon publication all high throughput sequencing data will be made available to the scientific community in the Sequence Read Archive (SRA) and Gene Expression Omnibus (GEO) databases. The plasticity scores calculated in large population studies are available as part of the Supplementary Information. All other data supporting the results of this study are available from the corresponding author on reasonable request.

## Code availability

The R function used to compute plasticity scores is available as part of the supplementary material.

