## Supplementary results, supplementary methods, Supplementary Figures, Supplementary Figures and Tables legends for "Identifying cancer cell-state transitions from multimodal single-cell data"

### Single-cell gene expression dynamics identify the molecular determinants of leukemia cell plasticity

#### Supplementary results

##### CRISPR-KO screening and data analysis pipeline validation

To validate our CRISPR-KO screening platform, we compared CD24+ and CD24- cells after the first sorting step (Figure 4A) to identify sgRNAs associated with the CD24+ phenotype (Table S7). Indeed, genes required for CD24 protein biosynthesis were expected to emerge among the phenotype-associated hits, providing an internal benchmark for the screen and analysis pipeline.

As expected, sgRNAs targeting CD24 and pathways required for CD24 biosynthesis, including GPI-anchor metabolism, were depleted in CD24+ populations (Figure S5A-B, Table S8). CD24+ cells also showed depletion of sgRNAs targeting genes related to mitochondrial homeostasis and oxidative metabolism, consistent with their distinct metabolic profile. Interestingly, several chromatin-remodeling factors, including RCOR1 and KDM1A, were among the top candidates, suggesting a potential role in stabilizing the CD24- phenotype (Table S7).

##### Rank deviation score validation

To validate our scoring method, we compared our rank-deviation-based plasticity scores to single-sample GSEA (1) using the genes overexpressed in the transitioning cells (N=46). In a cohort of 96 CML patients initiating Imatinib treatment (2), the two scores were highly correlated ($r^2 > 0.9$, $p < 2*10^{-16}$; Figure S6) supporting our scoring approach. Notably, our approach also accounts for the genes repressed in state transition (N=10) and allows to account for sampling site. Therefore, the rank deviation score was chosen for all the analyses presented.

##### Plasticity signature identifies proliferative hotspots in solid tumors

To investigate the spatial distribution of cells expressing the plasticity-associated program in solid tumors, we reanalyzed public spatial transcriptomics data from HCC patients (3). This dataset comprised two Visium sections, CHC20 and CHC23, from two different patients. Both sections were dominated by tumor cells but displayed distinct histopathological features. CHC20 contained a broad fibrotic region, whereas CHC23 included a large proliferative domain and a necrotic area characterized by a high mitochondrial transcript fraction (Figure S8A-B).

As summarized in the main Results section, we computed a plasticity score for each Visium spot and modeled its association with ssGSEA-derived Gene Ontology biological process scores, adjusting for sequencing depth, library complexity, and patient identity. Plasticity scores correlated positively with signatures related to proliferation, organelle division, and oxidative metabolism, and negatively with fatty acid metabolism programs (Table S17). Spatial mapping showed that high-plasticity spots formed discrete intratumoral niches. In CHC20, plasticity peaked within the fibrotic region, which also showed elevated proliferation, oxidative phosphorylation, and organelle fission scores (Figure 6C-F). In CHC23, plasticity was highest in a proliferative tumor domain enriched for analogous pathways (Figure S8C-F).

To assess whether spatially localized plasticity-associated programs were conserved in an independent solid tumor context, we analyzed spatial transcriptomic profiles from clear cell renal cell carcinoma (ccRCC), comprising 12 sections from 12 patients (4). As observed in HCC, the plasticity score correlated positively with oxidative metabolism, proliferation, and organelle-remodeling signature scores (Figure S9 and Table S18), supporting the presence of localized high-plasticity regions associated with proliferation-linked mitochondrial programs across distinct solid tumor types.

### Supplementary Materials and Methods

#### Public single-cell RNA-seq data re-analysis

All count data analyses were performed using Seurat (v5.3.0) (5) in an R v4.4.0 environment.

*K562 SMART-Seq dataset*

Single cell RNA-seq data were obtained from the Gene Expression Omnibus (GEO) database (GSE85534) (6). Gene expression quantification was performed using kallisto software (7) and the Gencode v40 human genome annotation. To reduce experimental bias, quality control was performed evaluating the number of reads per cell, the number of genes detected per cell, and the percentages of reads aligning to mitochondrial and ribosomal genes. Cells displaying poor data quality were removed from the analysis. Data normalization was performed using the Seurat package (5). Normalized counts were log-transformed and scaled, co-expression of genes with the CD24 was evaluated by Pearson correlation. Multiplicity of testing problem was addressed using the Benjamini-Hochberg false discovery rate. Adjusted p-values < 0.1 were considered statistically significant.

*Bone marrow CITE-Seq dataset*

Bone marrow CITE-seq data (8) were obtained from GEO (GSE245108). From the original study, we used the dataset generated after antibody titration, including patients BF21, BM27, WF26, and WM34. To avoid distortions in protein-level estimates due to sequential staining procedures with antibodies targeting the same antigens, only unsorted samples were included in the analysis.

Strict quality control was applied to maximize the signal-to-noise ratio. Briefly, cells with fewer than 1,500 detected RNA features, fewer than 3,000 RNA UMIs, fewer than 50 detected ADTs, fewer than 500 ADT UMIs, more than 20% mitochondrial RNA UMIs, or more than 15,000 ADT UMIs were removed. Only RNA features with at least 20 UMIs detected in at least 10 cells and ADTs with at least 5 UMIs detected in at least 3 cells were retained.

Doublets were removed by k-nearest-neighbor (kNN) analysis using the scDblFinder R package (9). To remove antibody aggregates, cells above the 99.95th percentile for each antibody were excluded. The final dataset included 18,920 cells, 23,721 RNA features, and 141 protein features.

The antibody matrix was normalized using the DSB method with approximate background distribution (10), whereas RNA features were normalized using the standard Seurat workflow. Cell classification against a bone marrow reference dataset was performed using BoneMarrowMap (11) with standard parameters and a maximum mean absolute deviation of 2.5.

For data visualization, Harmony reduction was used to remove interindividual variability. UMAP was performed on the Harmony reduction using 30 components and 50 nearest neighbors.

To select cells belonging to the monocyte differentiation path, we retained cells with broad classifications corresponding to hematopoietic stem cells or multipotent progenitors (HSC/MPP), lympho-myeloid primed progenitors (LMPP), cycling progenitors, early or late granulocyte-monocyte progenitors (GMP), pro-monocytes, or monocytes. From these, cells with narrow classifications indicating commitment to other lineages, including megakaryocyte-erythrocyte-primed MPP and neutrophil-primed GMPs, were excluded from the analysis. Because the analysis focused on classical monocyte differentiation occurring in the bone marrow, CD16 monocytes were also excluded. The final monocyte-lineage dataset comprised 5,341 cells.

UMAP was performed on Harmony reduction using 20 components and 30 nearest neighbors. Trajectory analysis was performed using Monocle 3 on the same UMAP embedding, with Leiden clustering and k = 10. The root for pseudotime ordering was set proximal to hematopoietic stem cells.

#### Cell metabolic profiling

K562 cells were sorted using a FACSAria III cell sorter (BD Biosciences) and an anti CD24-PE antibody (BD Biosciences, 560991). Briefly, cells were washed with PBS and incubated for 15' at +4°C with 5ng/µl CD24-PE antibody. Flow sorting was performed using a 100 µm nozzle and at controlled temperature. Sorted cells were recovered in Seahorse XF RPMI assay medium with 2mM L-glutamine, 10 mM glucose and 1mM pyruvate at pH 7.4 (Agilent, 103681-100). CD24+, CD24- and unsorted cells were seeded at a density of 112,500 cells per well in Cell-Tak (Corning, 354240) coated 96 well plates. Baseline and stressed energetic phenotype were assessed using Seahorse XF Cell Energy Phenotype

Test Kit (Agilent, 103325-100) according to manufacturer's instructions and a Seahorse XFe96 analyzer (Seahorse Bioscience, Agilent Technologies). Oligomycin and carbonyl cyanide 4-(trifluoromethoxy) phenylhydrazone (FCCP) were applied at final doses of 1 μM and 0.5 μM, respectively. After the analysis, proteins were extracted using M-PER lysis buffer (Thermo Fisher) and protein amount per well was assessed by Bradford protein assay (Bradford reagent, Sigma-Aldrich) and protein/well values were used for data normalization. The analysis was performed on five independently sorted lines. Each independently sorted sample was assessed at least in technical duplicates, and the average measure of the technical replicates was considered. Raw data analysis was performed using Seahorse Analytics software (Seahorse Bioscience, Agilent Technologies).

#### K562 cells immunomagnetic sorting and flow cytometry analysis

K562 cells were sorted into CD24+ and CD24- populations using CELLection Pan Mouse IgG Kit (Thermo Fisher, 11531D) and an anti-CD24 antibody at 2.5 μg/ml (BD 555426, clone ML5) according to manufacturer's instructions. The CD24+ and CD24- fractions were assessed at T0, 3 days and 6 days post-sorting by flow cytometry.
To stain K562 cells for surface CD24 expression, they were washed with cold PBS and resuspended at 1 million/ml in FACS solution (0.5% in BSA in PBS) and CD24-PE (BD Biosciences, 560991) antibody was added at a concentration of 5ng/μl according to manufacturer's instructions. Cells were then incubated on ice for 15', washed in PBS, fixed in 4% formaldehyde in PBS for 15' at RT, washed again in PBS, and resuspended in PBS. Cells were washed with PBS once again and stored in FACS solution until analysis on a BD LSRII cytometer. Resulting data were processed using FlowJo software.

#### 10x scRNA-seq and feature barcoding

Three independent K562 samples at different passages were stained with a panel of hashing antibodies (BioLegend, TotalSeq B) following the manufacturer's instructions. Specifically, cells were stained with anti-CD24 (Cat. #311145), anti-CD34 (Cat. #343539), anti-CD38 (Cat. #356639), a mouse IgG2aκ isotype control (Cat. #400291), a mouse IgG1κ isotype control (Cat. #400185), and a unique cell hashing antibody for each technical replicate (Anti-human Hashtag 1–3, Cat. #394631, #394633, and #394635). CD24, IgG2aκ, and hashing antibodies were used at 0.5 μg/100 μL, while the remaining antibodies were used at 1 μg/100 μL. Briefly, $1*10^6$ cells were harvested, washed once in PBS, and resuspended in 45 μL cold Cell Staining Buffer (CSB) (BioLegend, Cat. #420201). Fc receptor blocking was performed with 5 μL Human TruStain FcX (BioLegend, Cat. #422301) for 10 minutes at 4°C. Antibodies were pooled and diluted in 50 μL CSB, centrifuged at 14,000 × g for 10 minutes at 4°C to remove protein aggregates, and the clarified supernatant was added to the cells. Staining was performed for 30 minutes at +4°C, followed by three washes with cold CSB. Samples were resuspended in PBS and pooled. Cell number and viability were assessed prior to library preparation. All samples had a viability >90%, and 10,000 cells were used for sequencing. scRNA-seq and feature barcoding libraries were prepared using the Chromium Next GEM Single Cell 3′ Reagent Kits v3.1 with Feature Barcoding technology for Cell Surface Protein (10x Genomics, Manual CG000317 Rev D) according to the manufacturer's protocol. Quality control of cDNA and final libraries was performed using the High Sensitivity DNA Assay on an Agilent 2100 Bioanalyzer. Feature barcoding libraries were sequenced on a NovaSeq 6000 platform at a depth of 10,000 reads/cell (total: 105.7 million reads). scRNA-seq libraries were sequenced in two batches using a NovaSeq 6000 and a NovaSeq X platform, at a depth of $\approx 4*10^5$ reads per cell (total: 3.9 billion reads).

#### 10x scRNA-seq and feature barcoding data analysis

*Reads processing, data normalization, and cleaning, CD24+ and CD44+ cells classification*

Alignment and gene expression quantification were performed using CellRanger v7.1.0 (10x Genomics). The GRCh38-2020-A reference genome and a custom antibody reference were used for analysis. For the scRNA-seq libraries, >95% of bases were Q30, 96.6% of reads mapped, sequencing saturation was 63%, and 9,291 cells were detected. Count data analysis was performed using the Seurat package (v5.3.0) (5). To remove cells associated with immune complexes, cells in the top and bottom 0.5% of total antibody UMI counts were excluded. Technical replicates were identified based on the ratio of HTO UMI per cell to total antibody UMI. As shown in Figure S1A, HTO UMI count distributions were examined across replicates. Cells positive for different HTOs were considered inter-sample duplicates. No significant batch effects were observed between replicates (Figure S1B). Additional filtering retained only cells with ≥20,000 UMI/cell and ≥6,000 detected features/cell (Figure S1C). To remove low-quality or apoptotic cells, those with <12% ribosomal UMI or >10% mitochondrial UMI were excluded. Low-count features (fewer than 20 total UMIs or detected in fewer than 10 cells) were also removed. Intra-sample duplicates were further identified using k-nearest neighbor (kNN) analysis with the scDblFinder R package (9), using HTO-based intra-sample duplicates as training set (Figure S1D). After duplicate removal, 6,900 cells × 23,995 features remained for downstream analysis. For CD24 normalization, CD24 antibody oligo counts were regressed against its isotype control (IgG2aκ), and the resulting residuals were scaled and used as a proxy for CD24 protein abundance (Figure S1E). Cells were then classified into CD24+ and CD24- populations based on these normalized protein levels (Figure S1F), cells with z-score < 0.3 were classified as CD24- while cells with *Z*-score > 0.5 were considered CD24+. CD44+ cells were instead classified based on the CD44 mRNA levels, all cells with CD44 UMI count > 0 were classified as CD44+.

*Dimension reduction and pseudotime analysis*

Cells were normalized using SCTransform (12), and the top 3000 highly variable genes were selected for dimensionality reduction. Principal component analysis (PCA) was performed using the Seurat R package. Uniform Manifold Approximation and Projection (UMAP) was computed using the first 30 principal components, with 50 neighbors and 1,000 iterations specified in the Seurat workflow. Variation in principal component scores across different state transition phases was assessed using the Kruskal–Wallis test. Pseudotime trajectory analysis was conducted using Monocle 3 (v1.3.7)(13), using 30 principal components and learning a trajectory graph across the full dataset. Cells in the erythroid state were designated as the root for pseudotime ordering. Differences in pseudotime across state transition phases were evaluated using ordinal logistic regression.

*CD24+ and CD44+ cells markers identification*

To identify CD24+ and CD44+ marker genes, cells were classified based on protein or mRNA expression levels as described above. Count data were normalized using the standard Seurat workflow, and differential expression analysis was performed using Seurat with Benjamini–Hochberg (BH)-adjusted false discovery rate (FDR) correction. Genes expressed in fewer than 1% of cells in either population were excluded. Genes with an absolute $\log_2$(fold change) > 0.1 and an adjusted p-value < 0.1 were considered statistically significant. Gene set enrichment analysis (GSEA) was performed using the GO database via the ClusterProfiler R package and GO term redundancy was addressed using ClusterProfiler simplify method. Cell type profiling was conducted by overrepresentation analysis of genes overexpressed in $CD44^+$ cells, using the enrichR R package (14) and the Azimuth 2023 reference (15).

*Cell cycle analysis*

Count data were normalized using the standard Seurat workflow. Cell cycle analysis was performed using the cyclone function from the scran package (16), based on the built-in cell cycle marker genes. Statistical analysis of population frequencies across different cell classes was conducted using the chi-squared test.

*Transitioning cells identification and pseudobulk analyses*

Log-normalized and scaled CD24 mRNA UMI counts were regressed against CD24 protein levels. A second-order polynomial model (Figure S3A) was employed, as it significantly improved predictive accuracy over a linear model ($p < 2*10^{-16}$, analysis of deviance). Predicted CD24 protein levels were then used to stratify cells into CD24+ and CD24- populations using the same thresholds applied to the observed protein data. Cells with discordant predicted and observed protein classifications were labeled as transitioning (Figure 2B-C). Pseudobulk differential expression analysis was conducted using DESeq2 to identify markers of transitioning cells. For this analysis, cells classified as CD24-, CD24+, Transition-/+, or Transition+/- were included. A further low-count filter was also applied, retaining genes with counts >1 in at least 10 cells. Raw UMI counts were aggregated by class and biological replicate using Seurat, and differential expression analysis was performed in DESeq2, including CD24 status as a covariate. Overrepresentation analysis was performed against the Gene Ontology (GO) database using the ClusterProfiler package, with separate enrichment analyses for upregulated and downregulated genes. Genes and pathways with Benjamini–Hochberg adjusted p-values < 0.1 were considered statistically significant.

*BCR-ABL1 activity estimation*

Aggregate expression of transcriptional targets of the BCR-ABL1 pathway was calculated using Seurat gene module scoring implementation. Log-normalized and scaled counts were used and as BCR-ABL1 targets we used genes downregulated by Imatinib inhibitors in Ph+ cells described before (17).

*K562 cell classification*

Cells were classified by projection onto a bone marrow reference using BoneMarrowMap (11), with standard parameters and a maximum mean absolute deviation threshold of 2.5. Rare cells classified as hematopoietic stem cells (HSC, N = 1), cycling progenitors (N = 6), burst-forming unit-erythroid cells (BFU-E, N = 9), or colony-forming unit-erythroid cells (CFU-E, N = 54) were grouped into an "early precursors" category.

#### Genome-wide CRISPR-Cas9 knockout screening

*Screening line generation*

The two independent CRISPR screening lines were generated as previously described (18). Briefly, K562 cells were lentivirally transduced with pLenti-Cas9-T2A-Blast-BFP to express a codon optimized, WT SpCas9 flanked by two nuclear localization signals linked to a blasticidin-S-deaminase – mTagBFP fusion protein via a self-cleaving peptide (derived from lenti-dCAS9-VP64_Blast, a gift from Feng Zhang, Addgene #61425). Following blasticidin selection, a stable BFP+ population was isolated by repeatedly sorting for BFP expressors. The genome-wide Brunello sgRNA library (19) was synthesized as 79 bp long oligos (CustomArray, Genscript). The oligo pool was made double-stranded by PCR to include an A-U flip in the tracrRNA (20), 10 nucleotide long unique molecular identifiers (UMIs), and an i7 sequencing primer binding site (18). The resulting PCR product with the sequence was cloned by Gibson assembly into pLenti-Puro-AU-flip-3xBsmBI (18). The plasmid library was input sequenced to confirm representation and packaged into lentivirus in HEK-293T (ATCC, CRL-3216) using plasmids psPAX2 (a gift from Didier Trono, Addgene #12260) and pCMV-VSV-G (a gift from Bob Weinberg, Addgene #8454). The virus-containing supernatant was concentrated with Lenti-X concentrator (Takara), aliquoted and stored in liquid nitrogen. The functional titer of the library virus was estimated from the fraction of live cells after transduction of target cells with different amounts of virus and puromycin selection. Cas9-BFP-expressing target cells were transduced with the library virus in duplicate at an approximate MOI of 0.3 and a coverage of 1000x (1000 cells per guide) in the presence of 2 µg/ml polybrene. Transduced cells were selected with 2 µg/ml puromycin from day 2 to day 10 post transduction. Cell numbers per replicate were kept at >= 80 million/replicate throughout to ensure full library coverage. The cells were further cultured and expanded for 2 days without puromycin and then frozen for later use.

*Sequential sorting and library preparation*

Sorting procedure is outlined in Figure 3A. Two replicates were performed. Briefly, KO library cells (120 million/replicate) were thawed and plated in fresh culture medium. At day 3 all cells were pooled and 40 million cells were plated in fresh medium to be sequenced as unsorted control. The remaining cells (~250 million/replicate) were sorted into CD24+ and CD24- populations using CELLection Pan Mouse IgG Kit (Thermo Fisher, 11531D). All MACS sorting experiments were performed by staining the cells with an unlabeled anti-CD24 antibody at 2.5 µg/ml (BD 555426, clone ML5) and according to manufacturer's instructions. CD24+ and CD24- fractions (~10 and 180 million cells, respectively) were replated in fresh culture medium and cultured for 24 hours. After this time, part of the cells was collected for sequencing. 40 million cells were collected from the negative fraction, but given the lower number of cells, only 2 million cells were collected from the positive fractions. The remaining cells were sorted again as described above yielding the CD24-, Transition -/+, CD24+, and Transition+/- populations. After the MACS all the samples were collected, washed with PBS, and pellets were frozen at -20°C.

Library preparation was performed as previously described (18). Briefly, DNA was extracted using QIAamp DNA Mini kit (QIAGEN, 51304) according to manufacturer's instructions. DNA quantity and purity was assessed using a NanoDrop 2000c spectrophotometer (ThermoFisher). Sequencing libraries were prepared by running a series of three PCR reactions. Every reaction contained 0.02 U/µl KAPA HiFi HotStart polymerase (Roche, KK2502), 1X KAPA HiFi Fidelity buffer, 0.3 mM dNTPs, and 0.3 µM of each PCR primer. All reactions were performed as previously described (18). The resulting libraries were purified with 1.8X ratio of AMPure XP beads (Beckman Coulter, A63881) and sequenced on a NextSeq 2000 (Illumina).

#### *CRISPR-Cas9 knockout screening data analysis*

*Sequencing data preprocessing*

A custom data-mining pipeline was developed to identify gene-phenotype associations from two CRISPR-screening experiments. UMIs were extracted from raw reads using UMI-tools (21). Sequencing adapters and non-sgRNA sequences were trimmed using cutadapt software (22). Reads were aligned against the Brunello genome-wide library reference using bowtie2 (23). To address potential base-calling errors in UMI sequences, alignment files were grouped based on edit distances (≤1) using the UMI-tools directional grouping method with default parameters. Resulting tables were converted into an sgRNA UMI count matrix in R. Each sample was divided into 16 internal technical replicates based on the first two UMI sequence bases, as described previously (18). sgRNAs with extremely low counts (≤3 UMIs in fewer than five samples) were excluded from analysis. UMI counts were normalized using the distribution of control sgRNAs (N=1,000) and analyzed using DESeq2 as described below.

*CD24+ phenotype associated pathways*

For the CD24+ *versus* CD24- comparison, analyses were conducted on samples collected after the first sorting step. Differential sgRNA abundance was analyzed using DESeq2 adjusting for biological replicates. Only sgRNAs targeting genes expressed in K562 cells, confirmed by our scRNA-seq data, were retained. *p*-values were adjusted using the Benjamini-Yekutieli (BY) method to conservatively control the false discovery rate under arbitrary dependence among tests (24). sgRNAs with adjusted *p*-values < 0.1 were considered significant. Genes with discordant sgRNA associations were removed. To minimize biases from variable guide efficiency and to avoid overrepresentation of genes targeted by multiple sgRNAs in downstream pathway analyses, only the sgRNA showing the strongest statistical association per gene was retained for pathway-level enrichment. Gene Set Enrichment Analysis (GSEA) (25) was performed using the ClusterProfiler R package (26) and Gene Ontology (GO) database (27), with pathway-level *p*-values adjusted via the BH procedure.

*State transition associated pathways*

State transition-associated sgRNAs were identified using a strategy analogous to that employed for the CD24+ analysis and considering samples from the second MACS sorting step. Pairwise comparisons (-/+ transition *versus* stable CD24- cells and +/- transition *versus* stable CD24+ cells) and an overall comparison (all stable *versus* plastic cells) were conducted using DESeq2 (28). All analyses adjusted for biological replicate; the overall model was additionally adjusted for the initial cell state (CD24- or CD24+). Genes expressed in K562 cells were retained, and *p*-values were corrected for multiple comparisons using the BY procedure, adjusted p-values < 0.1 were considered statistically significant. sgRNAs showing significant associations in the same direction across the two pairwise and the overall comparisons were selected. Genes with discordant sgRNA associations were excluded from further analysis. Gene-level summaries and GSEA were performed as described in the paragraph above.

*Gene level analyses*

To better control for spurious or off-target effects of individual sgRNAs when investigating BCR-ABL1 in the CRISPR screen, all knockout events targeting the locus were considered. To identify BCR-ABL1-targeting sgRNAs, all BCR and ABL1 sgRNAs (N = 8) were aligned to the BCR-ABL1 fusion transcript sequence 9501_9501_1 from FusionGDB2, and sgRNAs with full-length alignments were selected (N = 3). UMI counts per gene were normalized with DESeq2 using control loci, and the association with plasticity was assessed by two-way aligned rank transform (ART) ANOVA accounting for state transition and initial cell state.

#### Clinical cohorts transcriptomics data analysis

*Plasticity score calculation and associations*

All analyses were performed using R v4.4.0. The detailed data preprocessing strategy for each cohort is described below. Normalized datasets were filtered to remove low-count features, and the plasticity signature score was derived as the signed sum of gene rank deviations from the average rank across the cohort. For each sample, genes were ranked by expression and compared to the average rank across all samples. Within each cohort, the score was calculated as the signed sum of these deviations, with the sign determined by the association direction in the original signature, and subsequently Z-score–normalized. Where specified, patients were stratified into high- and low-score groups based on the median score.

In cohorts including patients sampled in different sites (Beat AML, TARGET-AML) or different tumor types (TCGA pan-non-hematological-tumors) rank deviation, Z-score normalization, and patient stratification by plasticity score were computed by sampling site (AML cohorts) or tumor type (TCGA pan-non-hematological-tumors).

Single-sample GSEA (ssGSEA) (1) scores in the CML cohort were calculated using the GSVA package (29) with genes overexpressed in transitioning cells.

*Chronic Myeloid Leukemia (CML)*

Normalized gene expression data from 96 CML patients (2) were obtained from the Gene Expression Omnibus (GEO, GSE130404). In the original study, patients were profiled by microarray on peripheral blood (PB) at diagnosis and subsequently treated with Imatinib. All patients were in chronic phase, and early molecular response (EMR) at three months was used as a proxy for Imatinib resistance. For genes with multiple probes, the probe with the highest average expression was selected per gene.

Low-count features were excluded prior to analysis. Plasticity scores were computed using either the rank deviation method or single-sample GSEA (ssGSEA) (1). Associations between plasticity scores and drug response were assessed using Wilcoxon rank-sum test.

*Acute myeloid leukemia*

Gene expression and clinical data were obtained from the Oregon Health & Science University Beat AML study (30) via cBioPortal (31). Data from The Cancer Genome Atlas (TCGA-LAML) (32) and the TARGET initiative (33) were retrieved from the Genomic Data Commons (GDC) portal v43.0 (34). For Beat AML, we included patients profiled at initial diagnosis in peripheral blood (PB) or bone marrow (BM) with available gene expression and clinical annotations (N = 426). The TCGA-LAML cohort consisted of patients with primary AML profiled in PB (N = 142), and the TARGET cohort included patients with primary AML profiled in PB or BM (N = 1,907). TPM- or FPKM-normalized expression matrices were used as provided, with TPM preferred where available (TCGA, TARGET). Genes with mean expression <0.1 TPM or <0.5 FPKM were excluded. Plasticity scores were computed using the rank-deviation method as described above. Where indicated, cohorts were dichotomized into score-high and score-low groups based on the median. To minimize PB/BM-related bias, low-expression filtering, rank-deviation z-score calculation, and high/low class assignment were performed separately by sampling site. Associations between plasticity score and overall survival were assessed using Cox proportional hazards models adjusted for age.

*TCGA pan-non-hematological-tumors dataset*

Normalized gene expression and clinical data for TCGA pan-cancer samples (35, 36) were obtained from the UCSC Xena data hub (37), with additional clinical annotations retrieved from the GDC portal v43.0 (34) (initial N = 10,437). We included patients with primary, non-hematological (not leukemia or lymphoma) tumors who had both bulk tumor RNA-seq and clinical outcome data available and who had not received systemic therapy at diagnosis. To avoid incorporating samples misclassified in the legacy UCSC Xena dataset, patients absent from the most recent GDC release (v43) were excluded. In addition, TCGA GBM and LGG samples were reassigned into glioblastoma, oligodendroglioma, and astrocytoma according to recently proposed genomic criteria (38). The final analytic cohort comprised 8,873 patients. Genes with mean expression <0.1 TPM were excluded, and plasticity scores were computed using the rank-deviation method. To minimize bias introduced by tumor-specific expression patterns, low-expression filtering, rank-deviation z-score calculation, and high/low class assignment were performed by tumor type. Associations between plasticity score and available TCGA clinical

endpoints (36) were evaluated using Cox proportional hazards models adjusted for age. In pan-non-hematological-tumors analyses, tumor type was included as a stratification variable. In addition, sensitivity analyses were conducted using only tumor types with sufficient events to be reliably assessed for each clinical endpoint, as defined in the TCGA guidelines (36). To avoid bias introduced by the reclassification of glioblastoma (GBM) and low-grade glioma (LGG), all glioblastoma, oligodendroglioma, and astrocytoma cases were excluded from sensitivity analyses whenever either the original GBM or LGG cohort was deemed insufficiently powered for a given clinical endpoint.
Individual tumor type analyses were performed by age adjusted Cox regression. Cohorts with insufficient events to be reliably assessed for each clinical endpoint are clearly labelled in the forest plots.

##### Spatial transcriptomics data re-analysis

10x Visium data from two hepatocellular carcinoma (HCC) sections (3) and twelve clear cell renal cell carcinoma (ccRCC) sections (4), each derived from a distinct patient, were obtained from the Gene Expression Omnibus (GEO; accessions GSE245908 and GSE175540). In the HCC dataset, tissue morphology was evaluated using available high-resolution hematoxylin and eosin (H&E) images. In each dataset, spots were filtered based on library size (5,000 < UMIs < 100,000), complexity (2,500 < detected features < 10,000), and blood-derived RNA content (hemoglobin UMIs < 1%). Data were normalized using the SCTransform workflow (12), and per-spot plasticity scores were computed using the rank-deviation method. Gene sets corresponding to GO Biological Process terms were retrieved via the msigdbr R package, and ssGSEA scores were calculated per spot (1). To minimize patient-specific effects, plasticity and pathway scores were computed and Z-score–standardized independently for each slice. Associations between plasticity and individual pathways were modeled by linear regression, adjusting for sequencing depth, complexity, and slice of origin, and p-values were corrected for multiple testing using the Benjamini–Hochberg method. For visualization of selected signatures in spatial context, a nearest-neighbor smoothing approach was applied, whereby each spot was assigned the mean value of itself and its six nearest neighboring spots.

#### Supplementary Tables Legends

**Table S1.** Genes differentially expressed in CD24+ cells, as assessed by Wilcoxon test with Benjamini–Hochberg (BH) adjustment for multiple comparisons.
**Table S2**. Gene Ontology terms enriched among genes differentially expressed in CD24+ cells, as assessed by gene set enrichment analysis (GSEA).
**Table S3.** Genes differentially expressed in CD44+ cells, as assessed by Wilcoxon test with Benjamini–Hochberg (BH) adjustment for multiple comparisons.
**Table S4.** Cell type marker enrichment analysis (Azimuth database) using genes overexpressed in CD44+ cells.
**Table S5.** Genes associated with state transition in K562 cells, identified by DESeq2 differential expression analysis comparing transitioning versus stable cell populations.
**Table S6.** GO terms enriched among genes downregulated in transitioning cells, as assessed by overrepresentation analysis.
**Table S7.** sgRNAs enriched or depleted in CD24+ cells after the first round of sorting. Analysis performed using DESeq2.
**Table S8.** GO terms enriched in genes targeted by sgRNAs depleted or enriched in CD24+ cells, as assessed by gene set enrichment analysis (GSEA).
**Table S9.** sgRNAs enriched or depleted in transitioning cells. Results include pairwise comparisons (CD24- vs. Transition-/+, and CD24+ vs. Transition+/-, shown in blue and red respectively) and the overall model. All analyses were performed using DESeq2.
**Table S10.** GO terms enriched in genes targeted by sgRNAs depleted or enriched in transitioning cells, as assessed by gene set enrichment analysis (GSEA).
**Table S11.** Plasticity score values calculated for the Beat AML, TCGA-LAML, and TARGET AML cohorts.
**Table S12.** Cox proportional hazards regression results for overall survival across three AML cohorts: Beat AML, TARGET-AML, and TCGA-LAML, as well as the pooled AML cohort. Associations between the plasticity score, modeled either as a Z-score or as a binary high/low variable, and survival were adjusted for age.
**Table S13.** Plasticity score values calculated for the entire TCGA dataset.
**Table S14.** Multivariable Cox proportional hazards regression results for overall survival, progression-free interval, disease-free interval, and disease-specific survival across TCGA non-hematological cancer patients. Associations between the plasticity score, modeled as a Z-score, and clinical outcomes were adjusted for age, with tumor type included as a stratification variable.
**Table S15.** Multivariable Cox proportional hazards regression results for overall survival, progression-free interval, disease-free interval, and disease-specific survival across TCGA non-hematological cancer cohorts. Only tumor types with enough events to support estimation for each clinical endpoint, as assessed by the TCGA consortium, were included. Associations between the plasticity score (modeled as a Z-score) and clinical outcomes were adjusted for age, with tumor type incorporated as a stratification variable.
**Table S16.** Multivariable Cox proportional hazards models were fit for overall survival and progression-free interval separately within each TCGA non-hematological cancer type. Plasticity scores (expressed as Z-scores) were evaluated while adjusting for age. Reported values include hazard ratios (HR), 95% confidence intervals (CI), and *p*-values. Flags * and ** mark tumor types that TCGA guidelines identify as not recommended or recommended with caution for analyses of the corresponding endpoint.
**Table S17.** Summary statistics for associations between per-spot plasticity scores and GO Biological Process ssGSEA signatures across two HCC Visium sections. Reported metrics include the regression coefficient ($\beta$), standard error, *p*-value, and Benjamini–Hochberg–adjusted *p*-value. All models were adjusted for patient of origin, sequencing depth, and library complexity.
**Table S18.** Results of linear regression analyses testing the association between spot-level plasticity scores and GO Biological Process ssGSEA signatures across 12 ccRCC Visium sections. Reported parameters include the regression coefficient ($\beta$), standard error, *p*-value, and Benjamini–Hochberg–adjusted *p*-value. All models were adjusted for patient identity, sequencing depth, and library complexity.

#### Supplementary Figure Legends

**Figure S1. Read processing, data normalization, and cleaning.**

**A)** Single-cell RNA sequencing was performed on three independent K562 lines labeled using cell hashing antibodies conjugated with replicate-specific Hashtag Oligos (HTO). The figures show the HTO Unique Molecular Identifiers (UMI) counts per cell across the technical replicates. Cells with HTO signals from multiple samples were flagged as inter-sample duplicates. **B)** No clear batch effect was observed among the replicates. The panel shows the transcriptomic data dimension reduction by Uniform Manifold Approximation and Projection (UMAP) colored by technical replicate. **C)** The dataset was further filtered by removing cells with <20,000 UMIs or <6,000 genes. The plot shows the UMI counts per cell and the number of detected genes per cell. Cells were also filtered based on the ribosomal and mitochondrial UMI proportions, and low-count features were also removed (not shown). **D)** Intra-sample duplicates were also identified using scDblFinder and leveraging the HTO-based inter-sample duplicates as training set. The plot shows the UMI counts and detected gene distributions in intra-sample and inter-sample duplicates. **E)** CD24 antibody levels were normalized using its matched isotype control as a reference. Briefly, CD24 antibody oligo UMI counts were regressed against those of the IgG2aκ isotype controls. The scaled residuals were used as a proxy for the CD24 protein abundance. The plot shows the linear correlation between anti-CD24 antibody signal and IgG2aκ isotype control UMI distributions. **F)** The normalized protein levels were used to classify the cells in CD24+ and CD24-. Cells with Z-score < 0.3 were classified as CD24- while cells with Z-score > 0.5 were considered CD24+. The plot shows the normalized protein levels (*i.e.* scaled anti-CD24 antibody–IgG2aκ regression residuals) distribution in cells classified as CD24- and CD24+.

**Figure S2. CD24+ cell phenotype.**

**A)** UMAP projection of scRNA-seq data from K562 cells colored by cell classification after similarity analysis with bone marrow cell populations. EBL, erythroblast. **B)** Distribution of differentiation-stage classes in CD24+ and CD24- K562 cells. Statistical significance was assessed using a chi-squared test. **C)** Gene Ontology (GO) terms enriched in genes positively or negatively correlated with CD24 mRNA levels in publicly available K562 SMART-Seq data, as assessed by Gene Set Enrichment Analysis (GSEA). **D and E)** Basal Oxygen Consumption Rate (OCR, D) and Extracellular Acidification Rate (ECAR, E) in sorted K562 cells as assessed by extracellular flux analysis. The experiment was conducted on 5 independently sorted K562 samples and data are presented as mean ± standard error. Statistical analysis was performed by analysis of variance (ANOVA) and post-hoc Tukey Honestly Significant Difference test.

**Figure S3. Identification of transitioning K562 cells in scRNA-seq data.**

**A)** Second-order polynomial regression predicting CD24 protein levels from CD24 mRNA abundance. Cells are colored by protein expression class, as defined in the Supplementary Methods and Figure S1. **B)** Proportion of cells in each phenotypic class (CD24+, CD24-, Transition -/+, Transition +/-, and CD44+). **C-D)** Distributions of UMAP component 1 (C) and component 2 (D) coordinates by phenotypic class. **E)** Monocle 3 pseudotime distribution in K562 cells stratified by differentiation stage inferred using BoneMarrowMap. **F)** Pseudotime distribution in K562 cells stratified by phenotypic class. For C, D, and F, statistical analysis was performed using the Kruskal-Wallis test followed by Dunn's post-hoc test with Holm correction for multiple comparisons. CD44+ and unclassified cells were excluded from the analysis. For E, statistical analysis was performed using ordinal logistic regression, with differentiation stage as the dependent variable and pseudotime as the predictor.

**Figure S4. Canonical marker expression patterns during monocyte differentiation.**

**A)** UMAP projection of the complete public bone marrow CITE-seq dataset, colored by broad cell classification. **B)** UMAP projection of scRNA-seq data from bone marrow cells belonging to the monocyte differentiation lineage and selected for further analysis. HSC, hematopoietic stem cells; MPP, multipotent progenitors; LMPP, lympho-myeloid primed progenitors; GMP, granulocyte-monocyte progenitors; Mono, monocytes. **C)** Pseudotime trajectory analysis of the monocyte lineage. Black lines represent inferred cell-state trajectories, with numbered circles indicating the root point (white), branch points (black), and terminal states (grey).

**Figure S5. Genetic drivers of the CD24+ state.**
**A)** Volcano plot showing differential abundance of sgRNAs in CD24+ vs. CD24- cells after the first MACS sorting. Statistically significant sgRNAs selected for downstream analysis, as described in the Methods section, are marked in red. Briefly, sgRNAs with Benjamini–Yekutieli adjusted $p < 0.1$ were considered statistically significant, and genes targeted by sgRNAs with opposite effects were filtered out. S Selected sgRNAs are marked in red; filtered sgRNAs are marked in grey. **B)** Top 10 Gene Ontology (GO) terms enriched in genes targeted by sgRNAs depleted or enriched in CD24+ cells, as assessed by Gene Set Enrichment Analysis (GSEA).

**Figure S6. Validation of the plasticity scoring method.**
Correlation between rank-deviation–based plasticity scores and ssGSEA scores (based on genes upregulated in transitioning cells) in 96 CML patients investigated by microarray. Statistical analysis was performed by Pearson correlation.

**Figure S7. TCGA non-hematological cancers analysis.**
**A)** Forest plot showing associations between plasticity score and clinical outcomes across TCGA non-hematological cancers restricted to cohorts with robustly defined endpoints. **B)** Forest plot showing cohort-stratified associations between plasticity score and overall survival across TCGA non-hematological cancers. In **B**, * and ** denote cohorts not recommended or recommended with caution for this outcome, respectively, according to TCGA guidelines.

**Figure S8. Plasticity score in HCC spatial transcriptomics, CHC23 slice.**
**A)** H&E staining of the CHC23 section. Fibrotic regions and high cell density tumor areas are highlighted in black and yellow, respectively. No overt non-tumoral tissue was identified in this section. **B)** Spatial distribution of mitochondrial UMI fraction in CHC23. **C–F)** Spatial mapping of signature scores, smoothed using a mean nearest-neighbor method ($k = 6$), including plasticity (C), cyclin-dependent protein kinase activity (D), oxidative phosphorylation (E), and organelle fission (F).

**Figure S9. Plasticity score in ccRCC spatial transcriptomics.**
**A)** Top 20 GO Biological Process signatures most strongly associated with spot-level plasticity scores. **B–E)** Spatial mapping of Plasticity (B), cyclin-dependent protein kinase activity (C), oxidative phosphorylation (D), and organelle fission (E) in a representative ccRCC section. For visualization, signature scores were smoothed using a mean nearest-neighbor method ($k = 6$).

#### Supplementary Figures

**Figure S1. Read processing, data normalization, and cleaning.**

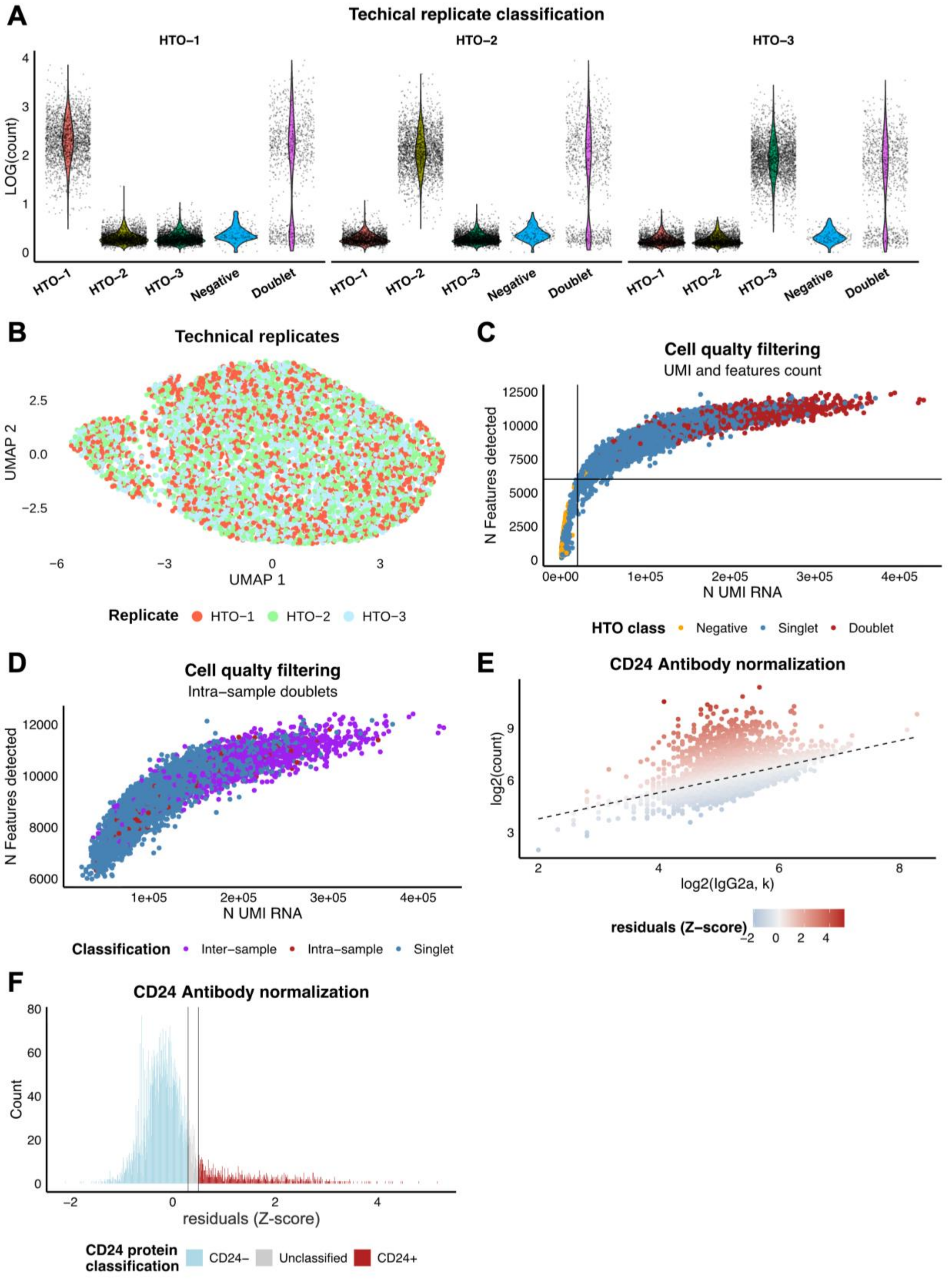

**Figure S2. CD24+ cell phenotype.**

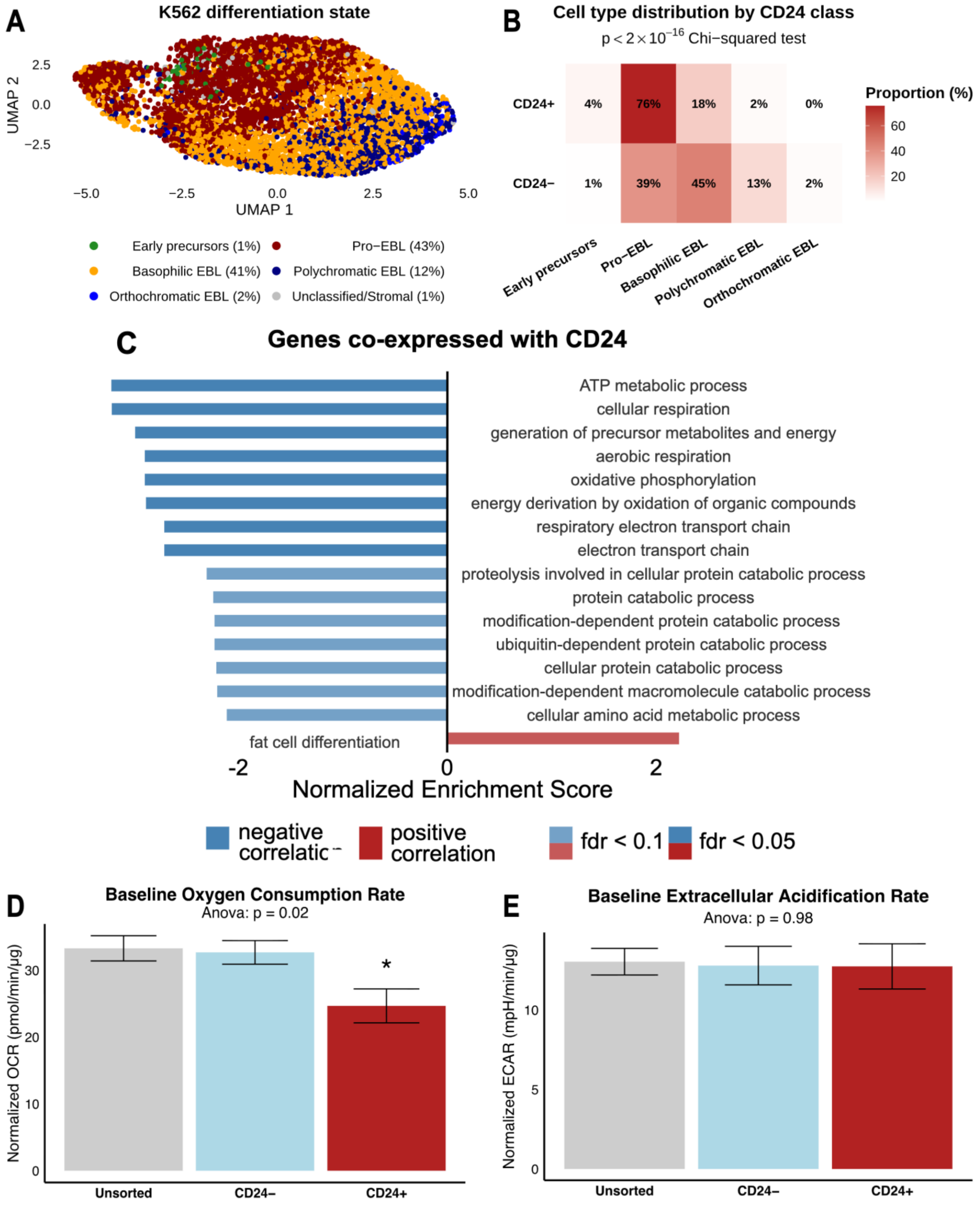

**Figure S3. Identification of transitioning K562 cells in scRNA-seq data.**

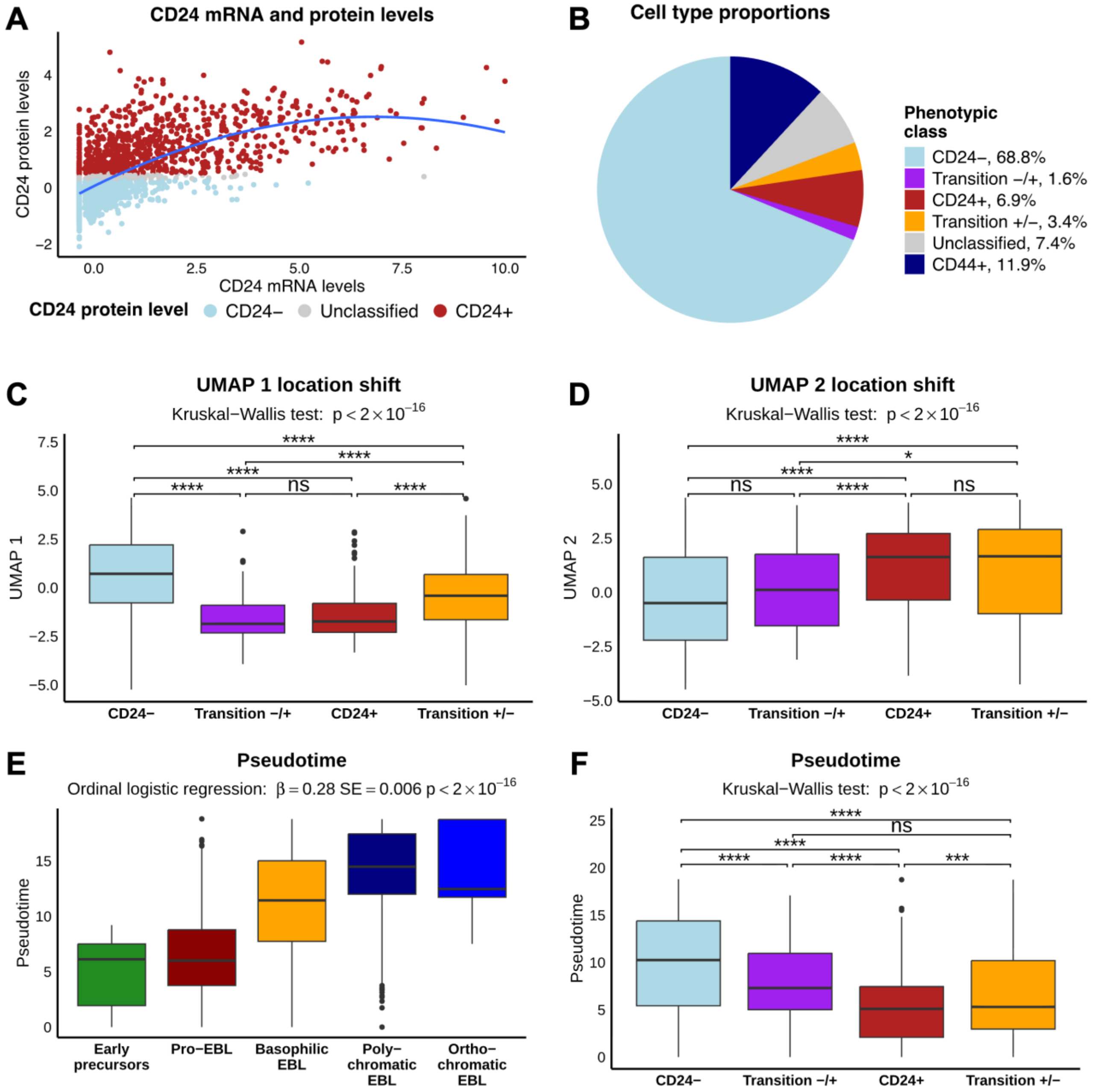

**Figure S4. Canonical marker expression patterns during monocyte differentiation.**

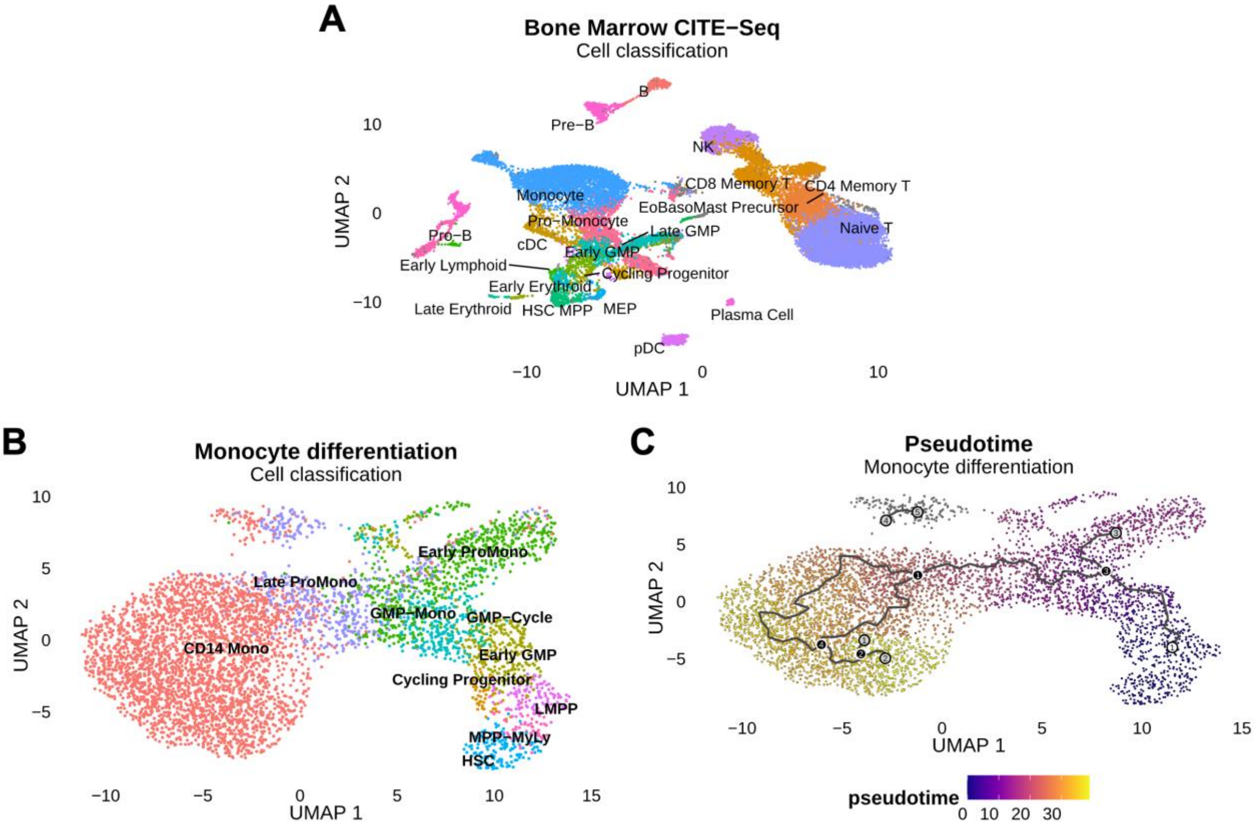

**Figure S5. Genetic drivers of the CD24+ state.**

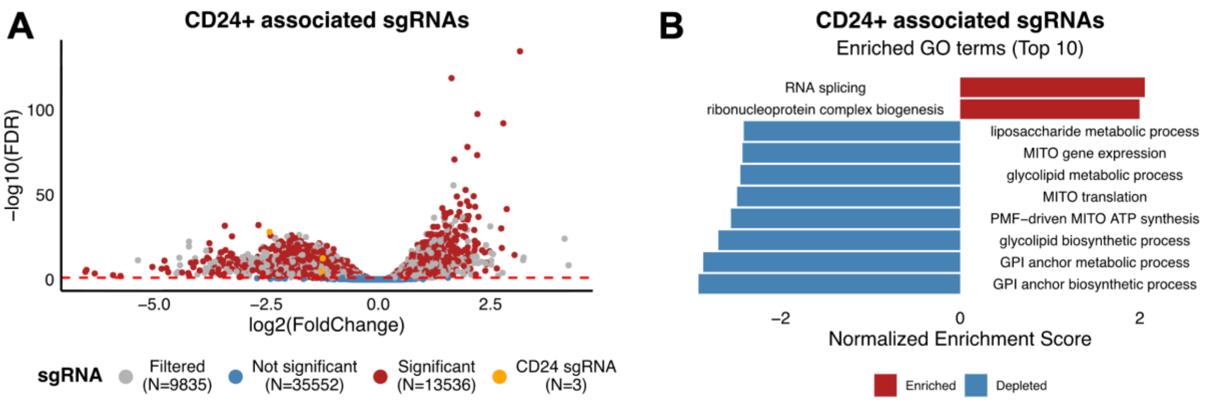

**Figure S6. Validation of the plasticity scoring method.**

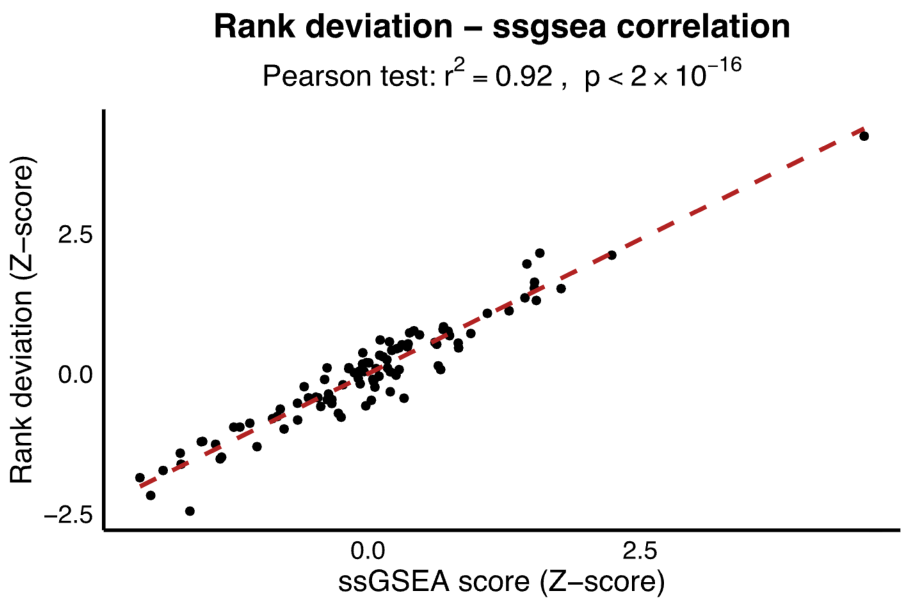

**Figure S7. TCGA non-hematological cancers analysis.**

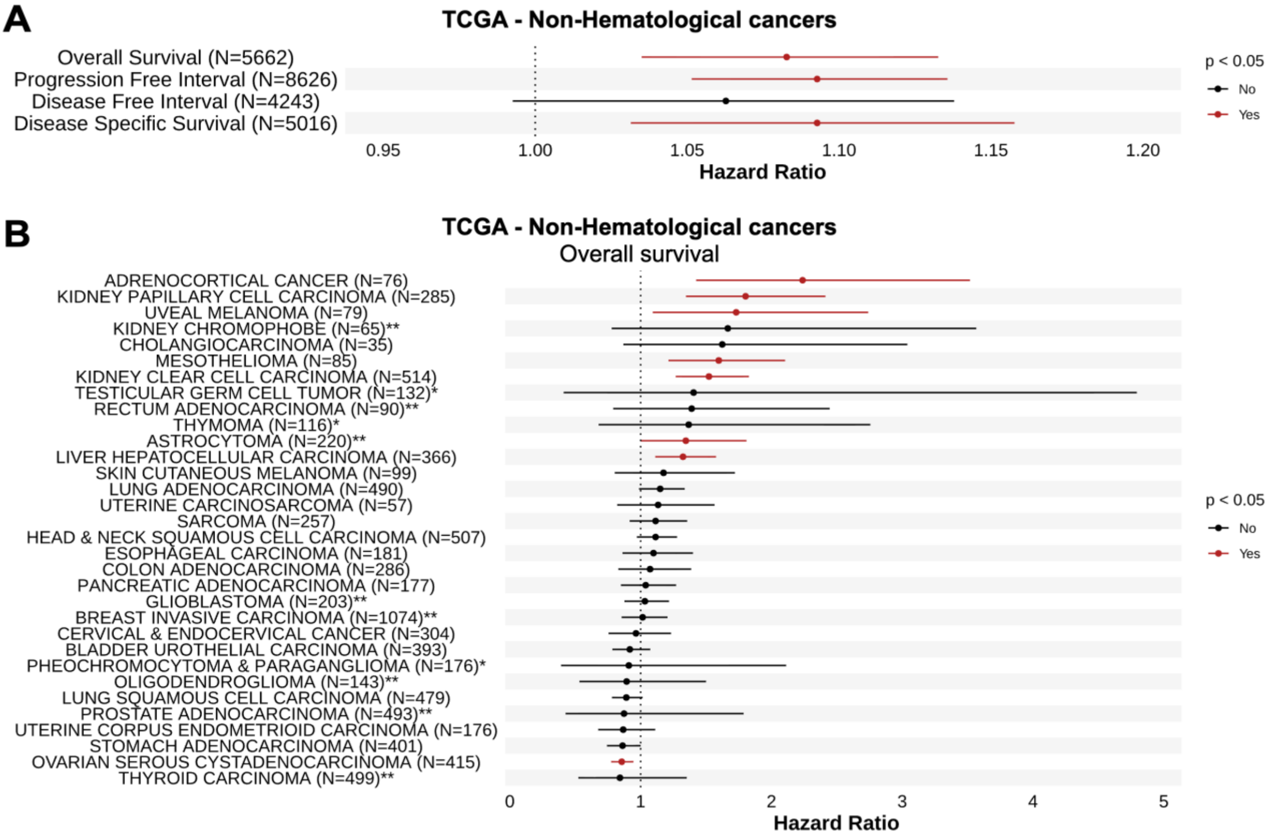

**Figure S8. Plasticity score in HCC spatial transcriptomics, CHC23 slice.**

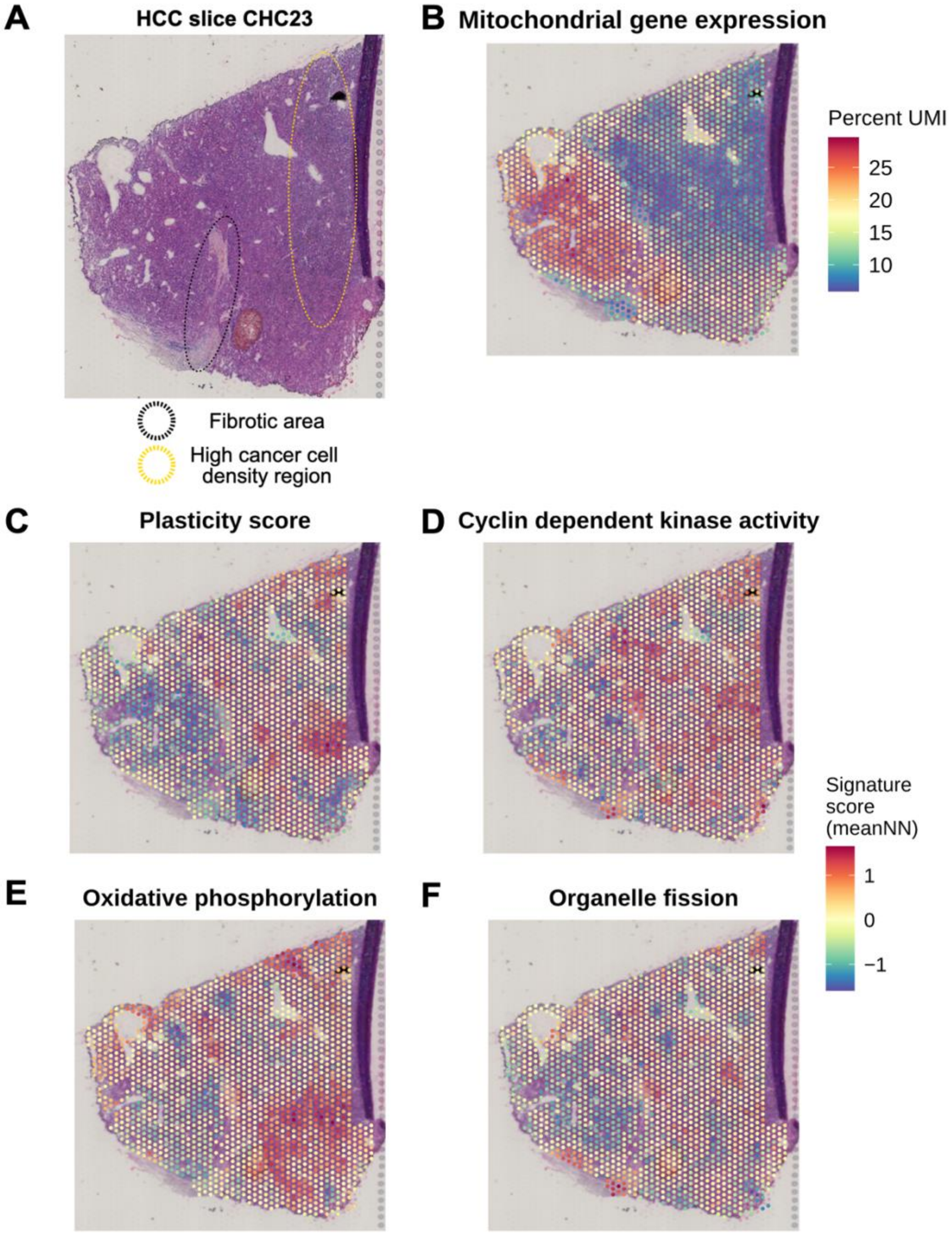

**Figure S9. Plasticity score in ccRCC spatial transcriptomics.**

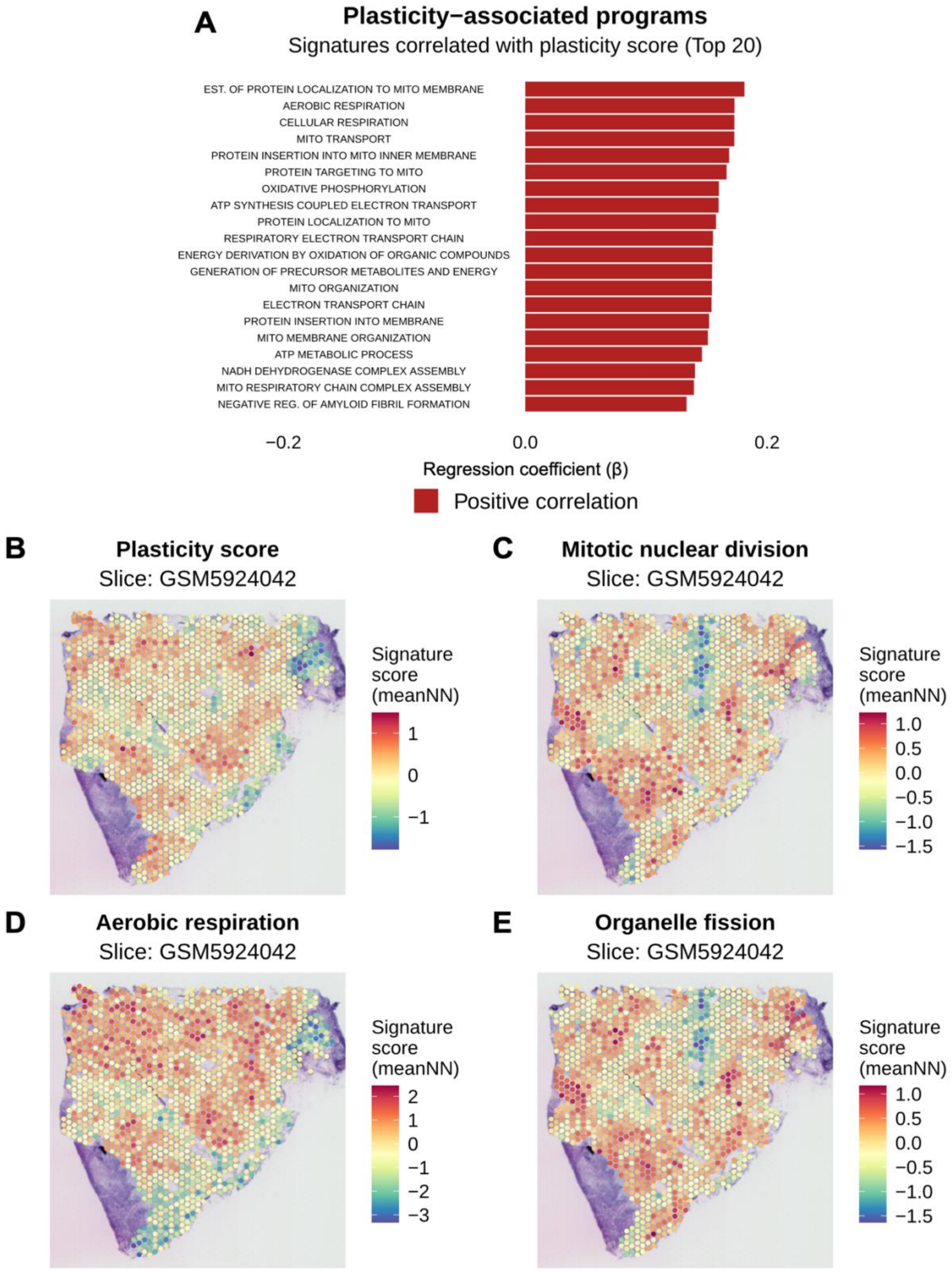
